# Extracting interpretable single-cell metabolic states with graph-guided representation learning

**DOI:** 10.64898/2026.09.17.751504

**Authors:** Daniel P. Lewinsohn, Nicolas Dias, Adelina Chau, Yuko Koike, Zachary D. Smith, Nilah M. Ioannidis, Allon Wagner

## Abstract

Metabolism shapes cellular function and state, yet measuring single-cell metabolic states at scale remains a challenge. We present Metabolic Representation Net (MeRN), a graph-guided variational autoencoder that leverages prior metabolic knowledge as a topology graph to learn latent representations of metabolic state and reaction activity from single-cell transcriptomes. MeRN’s scalable estimation of reaction activity enables the definition of *data-driven pathways* (DDPs): context-specific metabolic modules supported by transcriptomic evidence and agnostic of standard pathway definitions. Using DDPs, we introduce the *weakest link analysis* to identify metabolic network rewiring. MeRN recovers metabolic zonation in the mouse intestine, links a folate deficiency-induced break in *de novo* purine synthesis to embryonic neural tube defects, shows cytokines with similar non-metabolic effects can elicit divergent T cell metabolism, and identifies metabolic drivers of T cell exhaustion and therapy response in human cancers. Our results establish MeRN as a unified method for metabolic analysis of single-cell transcriptomes.

**Research highlights:**

- MeRN leverages the metabolic topology to comprehensively predict reaction- and pathway-level metabolic activities
- MeRN enables data-driven pathways (DDPs) that capture empirically supported cell-type-specific metabolic modules, agnostic of standard pathway definitions
- MeRN-based DDPs identify metabolic network rewiring of *de novo* purine synthesis due to folate deficiency during embryonic neural tube development
- MeRN identifies human pan-cancer metabolic drivers of T cell exhaustion and Treg-specific metabolic adaptations

## Introduction

Cellular metabolism is an essential aspect of cellular function and state, supporting the energetic and biosynthetic needs of the cell. Furthermore, dysregulation of cellular metabolic state, defined as the overall configuration of reaction fluxes and metabolite concentrations, is a central driver of many diseases, including cancer^1–3^, autoimmunity^4,5^, and neurodegenerative disease^6,7^. However, our ability to directly profile single-cell metabolic state with metabolomics remains limited by analytical sensitivity, robustness, metabolite coverage, and metabolite annotation confidence^8–10^.

An alternative method of measuring metabolic state utilizes single-cell transcriptomes. Single-cell RNA sequencing (scRNA-seq) broadly profiles the transcriptome, including enzyme-coding transcripts, henceforth referred to as metabolic genes, providing a comprehensive view of cellular metabolism. Importantly, scRNA-seq is standardized and scalable^11^, leading to its wide adoption and resulting in abundant datasets across all domains of biology, including translational applications to human health^12–16^. Additionally, scRNA-seq measures non-metabolic genes and can be paired with multimodal measurements of spatial location^17^, chromatin accessibility^18^, and genome-scale perturbation^19^, allowing metabolic states to be linked to other aspects of molecular cellular state.

Although promising, it remains difficult to comprehensively infer metabolic state from gene expression. Transcript abundance does not always correspond to enzyme concentration due to post-transcriptional and post-translational regulation^20^ and does not inform the directionality of reaction flux. Furthermore, existing gene-to-enzyme annotations are incomplete and do not fully account for many-to-many gene-reaction relationships. More generally, scRNA-seq suffers from technical noise, batch effects, and sparsity^21^. Finally, because metabolism operates as a highly interconnected biochemical network, deciding how to best map transcriptomic data onto this complex system remains an open problem.

Computational methods address these challenges by incorporating prior knowledge of metabolism into the inference of metabolic state from gene expression. Such methods can be broadly classified as constraint-based and pathway-based^22^. Constraint-based methods such as Compass^23^ and scFEA^24^ leverage the reaction-metabolite stoichiometric matrix as prior knowledge to infer reaction or module activity. While offering comprehensive inference of the metabolic topology, these methods scale poorly to large single-cell atlases while maintaining reaction-level resolution. Alternatively, pathway-based methods such as gene-set enrichment analysis (GSEA)^25^, overrepresentation analysis^26^, CellFie^27^, and scCellFie^28^ rely on predefined groupings of genes based on manually curated metabolic pathways. By pooling transcriptomic signal across the entire pathway, these methods leverage coordinated biological covariance to overcome technical sparsity and stochastic expression, enabling robust and highly scalable analyses. However, using predefined groupings of genes into pathways makes the strict assumption that all reactions in a given pathway will covary together in response to stimuli, which may not hold^29,30^. MetroSCREEN addresses this challenge by using gene-set enrichment for individual reactions, but it does not account for incomplete measurement of enzyme-coding genes across the metabolic reaction network^31^. Ultimately, the field lacks an approach that shares information across the metabolic network to reduce noise and account for missing enzyme measurements, while remaining agnostic to standard pathway boundaries within the graph.

In parallel, variational autoencoders (VAEs) have emerged as a primary method for analyzing single-cell transcriptomic data^32,33^. VAEs are highly scalable, can account for technical noise and batch effects, and produce information-rich cell embeddings. As a result, VAEs have been adapted for a wide range of tasks and applications in single-cell genomics^34–41^. More recently, these models have incorporated structured biological priors such as gene programs and molecular interaction graphs to learn interpretable cell embeddings or guide multimodal integration^42–44^, as did related latent-variable models^45^. Of particular interest is graph-linked unified embedding (GLUE), which jointly learns cell representations and embeddings of nodes in a prior regulatory graph, enabling integration across single-cell omic modalities^44^. The graph-guided VAE framework proposed by GLUE demonstrates how graph-structured prior knowledge can be incorporated into VAEs, providing a basis for integrating alternative graph representations, such as metabolic networks, to encode inductive biases.

Here, we present Metabolic Representation Net (MeRN), a graph-guided VAE for learning interpretable single-cell metabolic states. By jointly modeling single cells and a graph representation of the metabolic network, we enable comprehensive analysis and interpretation of metabolic state. The model simultaneously preserves reaction-level resolution while sharing information agnostically across reactions, thereby retaining the noise-reducing properties of pathway-based analysis without grouping reactions a priori.

MeRN offers three major advantages over previous methods. First, MeRN learns an interpretable metabolic latent space specifically capturing metabolic transcriptomic variation across cells, providing a powerful analytical tool for clustering, pseudotime analysis, and visualization. Second, the direct link between the metabolic latent space and the metabolic graph embedding enables *metabolic directions*: interpretable cell-type-specific metabolic state changes in response to stimuli or across time. Finally, MeRN’s flexible incorporation of prior knowledge enables the construction of metabolic *data-driven pathways* (DDPs). Although canonical pathways are indispensable for interpreting biological results, they may not reflect all physiological possibilities, especially in disease states. DDPs maintain the interpretive power of pathway analysis while offering evidence-based ties to mechanisms operating in specific biological contexts. Furthermore, DDPs enable analysis of metabolic network rewiring. We present MeRN-based DDPs as a novel framework for understanding cellular metabolism.

We demonstrate MeRN’s value in a variety of settings. First, we highlight our framework’s flexibility to model both reaction- and pathway-level activities across multiple metabolic axes in the small intestine, recovering known results and outperforming other methods in inferring metabolic states defined by protein expression – a molecular level closer to metabolic activity than mRNA expression. Second, we demonstrate that MeRN-based DDPs capture functional subunits of metabolism in the small intestine. Third, we use MeRN to show that folate deficiency during development rewires the metabolic topology of gastrulation, providing a mechanistic explanation for why dietary deficiency in an essential metabolite produces a highly specific embryonic neural tube defect. Fourth, we leverage MeRN’s interpretable latent space to disentangle metabolic and non-metabolic effects of cytokine perturbation in immune cells. Finally, we apply MeRN to a pan-cancer human T cell atlas, using DDPs to identify metabolic stages of CD8+ T cell exhaustion, recover known and unknown regulatory T cell (Treg) metabolic adaptations to the tumor microenvironment (TME), and associate the metabolic states of T cell subtypes with patient response to anti-PD-1 treatment in non-small cell lung cancer (NSCLC). These varied examples showcase the power of our graph-guided VAE approach to learning single-cell metabolic states.

## Results

### MeRN learns interpretable representations of single-cell metabolic state

MeRN is a graph-guided VAE that enables highly interpretable, exploratory analysis of single-cell metabolic states from scRNA-seq data. Inspired by previous work, we model cell states as low-dimensional cell embeddings learned by VAEs^32,46^, and we leverage prior biological knowledge in the form of a graph to guide our learned embedding^44^. Here, we model the transcriptomic profile of cells as complementary metabolic and non-metabolic low-dimensional embeddings **(Figure 1A)**. The metabolic embedding (M) reconstructs metabolic genes, and the non-metabolic embedding (B) reconstructs the remaining genes, thereby encouraging each latent space to capture distinct sources of variation.

**Figure 1.**
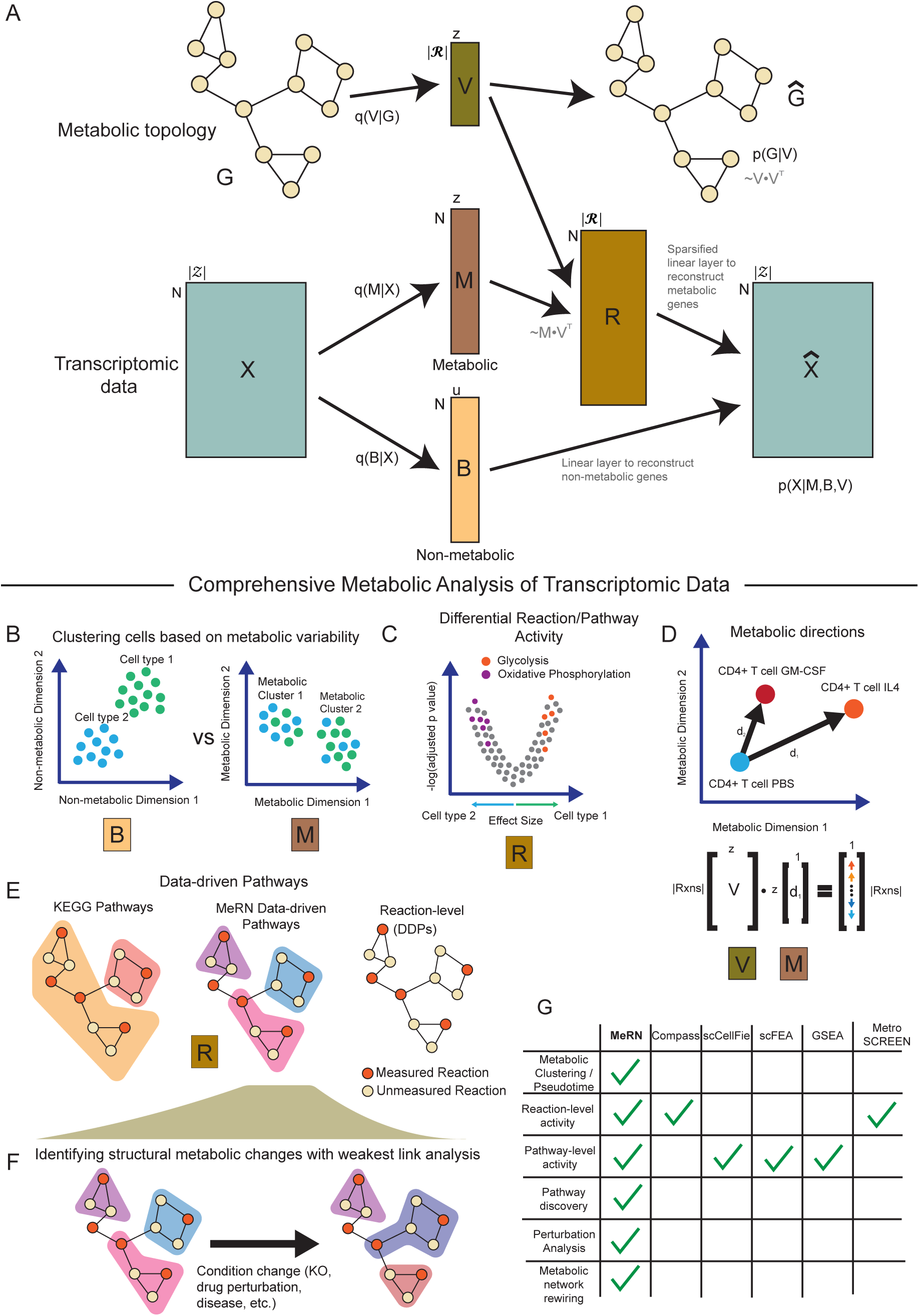
Learning interpretable single-cell metabolic states with graph-guided variational inference. **(A)** Illustration of MeRN architecture. MeRN inputs scRNA-seq data (X) and a KEGG-based graph representation of the metabolic topology (G). Neural network encoders (*q(V | G)*, *q(M | X)*, and *q(B | X)*) learn lower-dimensional embeddings for reactions (V) and cells, separated into metabolic (M) and non-metabolic (B) latent spaces. M is multiplied by *Vᵀ* to obtain the reaction activity matrix (R). R is decoded to reconstruct metabolic gene expression via a sparsified linear layer connecting reactions to transcripts encoding the enzymes for those reactions. Non-metabolic genes are decoded by a linear layer from B to *X̂*. Metabolic graph edges are decoded from pairwise dot products of the reaction embeddings in V. *p(G | V)* and *p(X | M, B, V)* denote the graph and scRNA-seq likelihood components. N denotes the number of input cells, z denotes the shared dimensionality of M and V, and u denotes the dimensionality of B. Throughout panels B-E, the labels M, B, and R indicate where model components are used in a typical analysis. **(B)** Cells can be clustered in either the metabolic or non-metabolic latent spaces, capturing distinct sources of biological variation. Each cell is denoted by a dot and is colored by cell type. **(C)** The reaction activity matrix can be used for differential activity analysis. Dots represent individual reactions. Blue and green correspond to cell types from (B). **(D)** The link between the metabolic and reaction latent spaces enables interpretation of directions in the metabolic latent space with respect to reaction changes. Dots represent the average location of different cytokine-perturbed states of CD4+ T cells. Direction vectors (*d₁* and *d₂*) can be calculated and projected through V to investigate reaction changes following perturbation. **(E)** MeRN-based data-driven pathways (DDPs) offer an alternative to predefined pathway- or reaction-level analyses. Nodes represent reactions, and edges connect reactions that share metabolites. Orange nodes represent reactions with transcriptomically measured enzymes; pale nodes represent reactions without such measurements (because of low expression or no matched enzyme). Colored halos group reactions by metabolic pathway. **(F)** The weakest link analysis with DDPs enables identification of metabolic network rewiring between reference (left) and perturbed (right) conditions. **(G)** Comparison of methods for metabolic inference. A check indicates that a method has explicitly demonstrated the corresponding capability.

To make the metabolic embedding interpretable, we encode metabolism as an undirected graph where nodes represent reactions and edges connect reactions that share metabolites. We construct this graph based on KEGG due to its extensive manual curation of experimental knowledge^47^ **(Methods)**. MeRN learns a low-dimensional embedding for each reaction included in the metabolic graph that captures the biochemical relationship between reactions. To interpretably link the metabolic embedding to the metabolic graph, we implement a specialized decoder that computes cell-specific reaction activity scores as the inner product of each cell’s metabolic embedding and the learned reaction embeddings. This computation results in a reaction activity matrix (R) **(Figure 1A)**, which captures the relative activity of every reaction across cells while effectively sharing information between reactions in the metabolic network. The reaction activity matrix then reconstructs the expression of metabolic genes through a sparsified linear layer, where reactions can only contribute to genes encoding an enzyme for the reaction, effectively limiting the metabolic latent space to only reconstruct metabolic genes. Reciprocally, the non-metabolic latent space is only allowed to reconstruct non-metabolic genes via a linear layer, limiting leakage between the latent spaces and enabling joint interpretation of non-metabolic gene programs^41^. We optimize all model parameters with mini-batch gradient descent to ensure scalability to large scRNA-seq datasets and perform hyperparameter optimization with a set of interpretability-based metrics **(Figure S1A-J)**.

Once trained, MeRN becomes a unified tool for analyzing single-cell metabolic states. The contrasting metabolic and non-metabolic latent spaces can be used for clustering, visualization, and pseudotime analysis **(Figure 1B)**, while the reaction activity matrix can be used for differential reaction or pathway activity analysis **(Figure 1C)**. To understand how cellular metabolism changes across relevant covariates (such as perturbation or time), we introduce *metabolic directions,* which describe how each reaction changes and can be used for clustering of metabolic and non-metabolic perturbation effects **(Figure 1D)**. Finally, we introduce DDPs as an alternative to predefined pathway- or reaction-level analyses **(Figure 1E)**. These can be used as a drop-in replacement for differential reaction or pathway activity analysis in a manner that improves interpretability without compromising their ability to explain nuanced metabolic variation. They can also be used to explore rewiring of the metabolic network between conditions via a *weakest link* analysis **(Figure 1F)**. MeRN’s versatility is unique among existing methods for metabolic state inference from transcriptomic data **(Figure 1G)**.

### MeRN resolves multi-dimensional metabolic zonation of enterocytes at pathway- and reaction-level resolutions

We applied MeRN to a previously generated scRNA-seq dataset of the mouse small intestine^48^ to test its ability to describe previously characterized metabolic variation of enterocytes at the single-cell level. Enterocytes are the primary absorptive epithelial cells of the small and large intestines^49^. Although enterocytes share a transcriptional identity, their metabolism varies significantly across the proximal-distal axis of the small intestine, as well as along individual villi **(Figure 2A)**^20,48^.

**Figure 2.**
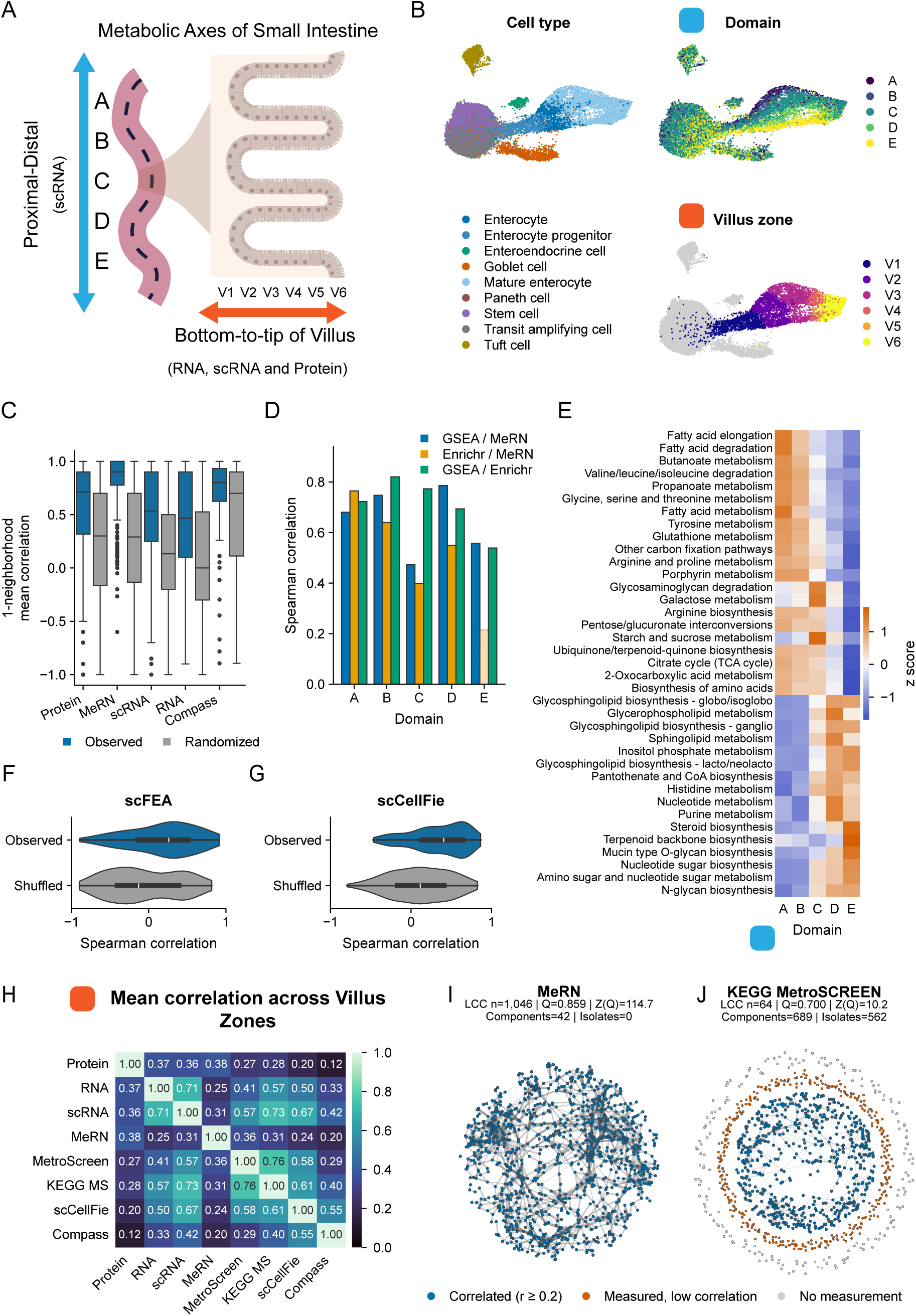
MeRN unifies pathway- and reaction-level metabolic analysis in the small intestine. **(A)** Enterocytes vary metabolically along the proximal-distal axis (stomach to large intestine; light blue) and from bottom to tip of individual villi lining the entire small intestine (orange). Analyzed datasets included scRNA-seq along the proximal-distal axis and scRNA-seq, bulk RNA-seq, and proteomics along the villus axis. Throughout this figure, panels addressing the proximal-distal axis are denoted by a light-blue square, and panels addressing the villus axis are denoted by an orange square. **(B)** Uniform Manifold Approximation and Projection (UMAP) representations of scRNA-seq data from the metabolic latent space, colored by cell type, proximal-distal domain, and villus zone. **(C)** For each reaction, the mean Spearman correlation between itself and its immediate neighbors in the metabolic topology network was computed across the five villus zones for each modality. Blue denotes the observed distribution, whereas gray denotes the corresponding distribution after 10 random relabelings of graph nodes. **(D)** Pathway activity was summarized in three ways: the one-versus-rest Cohen’s d of MeRN pathway activity (defined as the mean activity across reactions in each pathway), Enrichr combined score^92^, and GSEA normalized enrichment score (NES)^25^. Pathway summary scores were computed for each KEGG pathway within each proximal-distal domain. Spearman correlations were calculated among the MeRN, Enrichr, and GSEA pathway summary scores separately for each domain. Opaque bars indicate Benjamini-Hochberg-adjusted p < 0.05 across the 15 displayed tests; translucent, dashed bars are not significant. **(E)** KEGG pathway activity for each proximal-distal domain was computed by averaging reaction activity and calculating the z score across domains. The seven pathways with the largest MeRN Cohen’s d for each domain were selected for display. **(F)** Distribution of Spearman correlations across enterocytes between MeRN and scFEA scores for scFEA metabolic modules. For the shuffled baseline, scFEA module labels were permuted before correlation with the corresponding MeRN module scores; gray denotes shuffled correlations. **(G)** Same as (F), except comparing MeRN scores and scCellFie scores on the scCellFie-defined metabolic tasks. **(H)** Mean pairwise Spearman correlations between modality or method reaction profiles across villus regions, averaged across reactions shared between modalities or methods. **(I)** Force-directed layout of metabolic topology where nodes are reactions, and edges connect reactions that share a metabolite. For each retained edge, the weight is the Spearman correlation of MeRN activity between its two reactions across enterocytes; edges with correlations below 0.2 were omitted. **(J)** Same as panel (I), using MetroSCREEN reaction-activity correlations.

Zwick et al. characterized the metabolic zonation of enterocytes along the proximal-distal axis by segmenting and labeling the small intestine into 30 regions. The authors defined five distinct metabolic domains of enterocytes by clustering regions based on highly zonated genes. Visualizing the MeRN metabolic latent space with UMAP confirms that the metabolic latent space separates these five domains, without supervision or prior knowledge of cell state identities **(Figure 2B)**. We also confirmed this observation by computing the average position of enterocytes from each region in the metabolic latent space, calculating the pairwise Euclidean distances between all regions and performing hierarchical clustering **(Figure S2B)**. These results demonstrate that MeRN’s metabolic latent space captures metabolic variation of enterocytes along the proximal-distal axis of the small intestine.

To test whether MeRN describes the metabolic zonation of enterocytes along individual villi (bottom to tip), we leveraged a previous study by Harnik et al.^20^, who sorted enterocytes from the small intestine into six bins and performed bulk RNA-seq and proteomics. We constructed gene sets of differentially expressed genes for each region of the villi based on the bulk RNA-seq data. We then mapped these regionalized gene sets onto the scRNA-seq data from Zwick et al. by calculating a per-cell gene score **(Methods)**. Visualizing these gene scores on the metabolic latent space reveals regionalized expression of these gene sets along an orthogonal axis to the proximal-distal zonation **(Figure S2C)**. To assign a bottom-tip region to each enterocyte in the scRNA-seq dataset, we calculated pseudotime starting from V1 (bottom-most) region enterocytes **(Figure S2D, Methods)**. We then assigned each cell to a villus region based on which scaled region score was highest at a cell’s pseudotime **(Figure 2B, Figure S2E)**.

We leveraged this multimodal alignment of enterocyte profiling along the villus, including protein, bulk RNA-seq, and scRNA-seq, to validate one of the fundamental assumptions of MeRN’s design. Specifically, we assume the measured abundance of enzymes catalyzing biochemically related reactions in the KEGG metabolic network should be correlated. Indeed, immediate neighbors of each reaction are more correlated than a shuffled baseline across the villus in all three modalities, with MeRN recapitulating this trend **(Figure 2C)**. The result holds even when excluding immediate neighbors (reactions) that share a gene **(Figure S2F)**.

MeRN’s flexible graph-based representation of metabolism enables one to readily estimate arbitrary groupings of reaction activities, including predefined metabolic pathways. We tested these scores in the context of enterocyte zonation along the proximal-distal axis, beginning at the resolution of KEGG pathways. Across domains, MeRN, GSEA, and overrepresentation analysis pathway scores were significantly positively correlated, except for MeRN and overrepresentation analysis scores in domain E. These results suggest MeRN’s pathway scores are concordant with common pathway-based tools **(Figure 2D)**. To further assess the interpretability of MeRN pathway scores, we identified the pathways preferentially associated with each proximal-distal domain **(Figure 2E)**. We find that MeRN identifies strong zonation of many pathways in alignment with previous discoveries. For example, fatty acid metabolism localizes proximally to domains A and B^48,50^, as do components of the urea cycle (contained in KEGG’s arginine biosynthesis pathway)^48,51^. Starch and sucrose metabolism and galactose metabolism, components of dietary carbohydrate metabolism, are highest in domain C^48,52^, while steroid biosynthesis peaks distally in domain E^53^. Therefore, we conclude that MeRN identifies previously defined zonation of metabolic pathways along the enteric proximal-distal axis.

To further test MeRN’s flexible metabolic representation, we compared our single-cell estimations of metabolic activity to those from scFEA and scCellFie. Fundamentally, both tools group reactions into predefined pathways (metabolic tasks in scCellFie and modules in scFEA), with scCellFie following an approach akin to GSEA and scFEA using a flux-based approach. For each method, we calculated analogous single-cell MeRN metabolic task and module scores by averaging over reactions included in the pathways. Both methods showed strong correlations with MeRN against a shuffled baseline across enterocytes, further supporting MeRN’s ability to accurately estimate metabolism at many resolutions and bridge state-of-the-art modeling approaches **(Figure 2F,G)**.

Although still an incomplete picture of metabolism, protein measurement of enzymes is a stronger proxy for reaction activity than transcriptomic data as it bypasses the post-transcriptional and translational regulation layers between RNA and protein. In fact, Harnik et al. observed significant discordance between RNA and protein measurements along the villus, revealing that RNA concentration generally peaks closer to the bottom of the villi than protein concentration. This motivated us to leverage the paired protein measurements along the villus to assess the agreement and strength of different methods for inferring reaction-level activity.

To benchmark MeRN’s ability to predict reaction activity (as measured by protein concentration), we calculated gene expression profiles for enterocytes over the five regions of the villus where we were able to map to the single-cell data **(Methods)**. Next, we estimated the activity of each KEGG reaction with all modalities (protein, RNA, scRNA) and metabolic prediction methods (MeRN, Compass, MetroSCREEN, scCellFie) **(Figure S2G, Methods)**. We then calculated the average pairwise Spearman correlations for reactions with significant variation across the villus **(Figure 2H)**, showing the relationship between different prediction methods and modalities. As expected, we found strong concordance between reaction activity inferred by bulk and single-cell RNA, validating our mapping of villus zones from bulk to single-cell data. MeRN showed the highest correlation with protein measurements of reaction activity, followed by bulk RNA and scRNA measurements. Other methods such as MetroSCREEN, scCellFie, and Compass were highly correlated with each other and RNA measurements but did not outperform raw RNA measurements in predicting the protein measurements. These results further support MeRN’s ability to accurately estimate single-cell metabolic activity down to individual reactions across the entire metabolic network.

We next investigated MeRN’s global metabolic representation of enterocyte metabolism. Starting from the KEGG metabolic network used as MeRN’s input, we weighted undirected edges between reactions based on the Spearman correlation across enterocytes. We then removed edges with a correlation of less than 0.2 and visualized the network with the force-directed embedding algorithm **(Figure 2I)**^54^. We also visualized the metabolic network from MetroSCREEN adapted to KEGG **(Figure 2J)**. The graph produced by MeRN was more connected with 42 components and no isolates, whereas the MetroSCREEN graph had 689 components and 562 isolates (190 reactions not estimated and 372 reactions with no neighbors over the correlation threshold). Additionally, the largest connected component in the MeRN network contained 1,046 reactions and had a z-scored modularity of 114.7, while the MetroSCREEN network had a largest connected component with only 64 reactions and a z-scored modularity of 10.2 **(Figure S2H)**^55,56^. These results suggest that MeRN produces a substantially less fragmented, more modular, and more complete metabolic network representation of enterocytes compared to MetroSCREEN. This analysis prompted further exploration of clustering reactions in a data-driven manner based on MeRN’s metabolic estimations.

### MeRN enables data-driven metabolic pathway discovery

Predefined metabolic pathways represent a canonical partition of a complex metabolic network into discrete segments, and as such, may not capture functional subunits of metabolism most relevant for a given cell type, experimental condition, or disease. We therefore sought to identify groups of reactions which covary and are biochemically connected^57,58^. MeRN’s network-wide estimation of reaction activity enables partitioning of the fixed metabolic network topology into context-specific and empirically supported modules **(Figure 1E).** We term these modules *data-driven pathways* (DDPs), which we extract by performing a metabolic topology-constrained agglomerative clustering procedure based on reaction correlation across selected cells in the reaction activity matrix **(Figure 3A, Methods)**. We define three main applications for DDPs: (1) determining covarying, functional subunits of metabolism in a given cell type or state; (2) comparing usage of these functional subunits across cell types and experimental conditions; and (3) determining structural changes in DDPs between conditions or cell types to identify cases of metabolic network rewiring.

**Figure 3.**
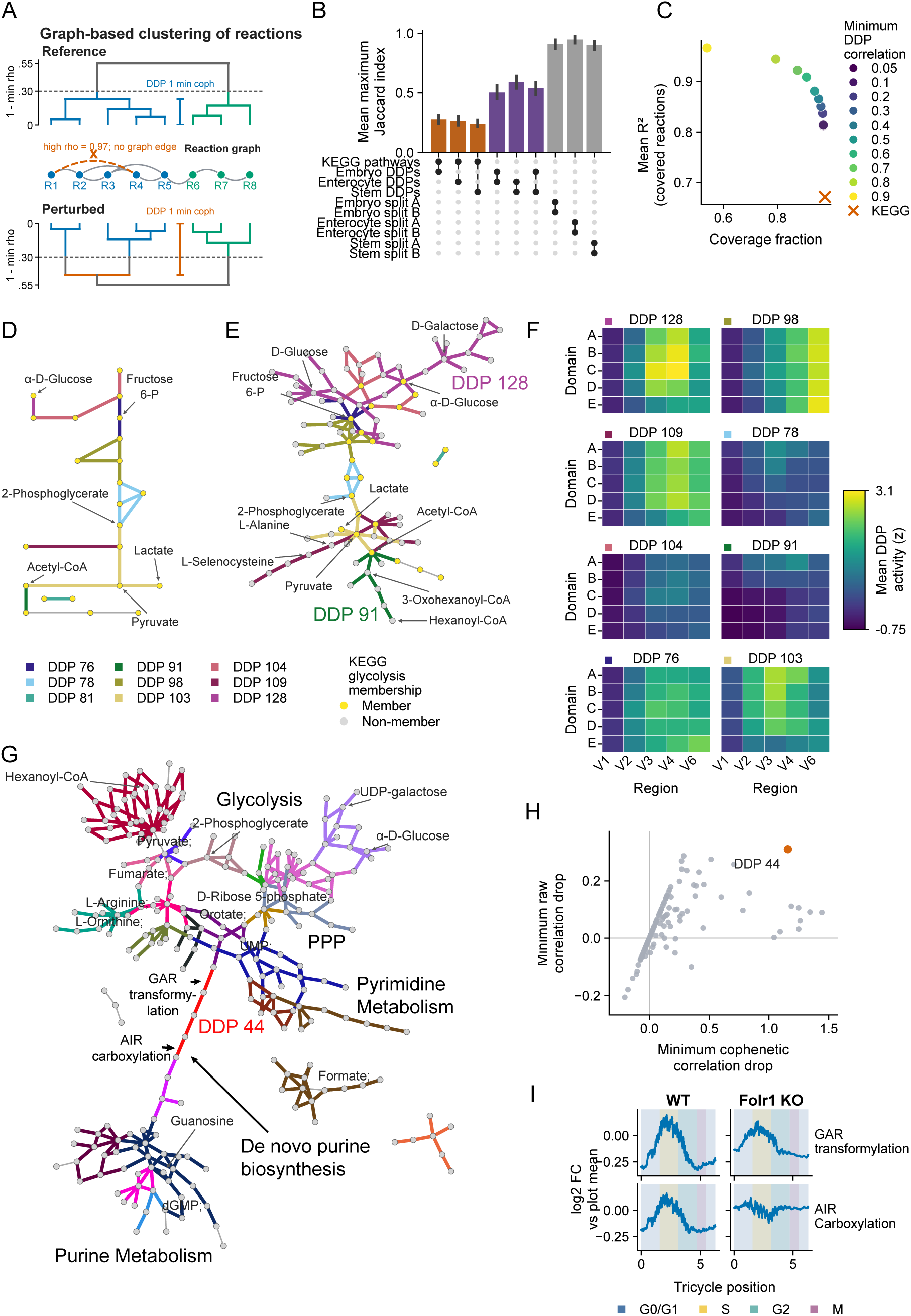
DDPs capture complex metabolic variation and reveal structural metabolic changes. **(A)** Conceptual diagram of DDP calculation under reference and perturbed conditions. DDPs are calculated via graph-constrained agglomerative clustering with complete linkage, meaning only reactions that are connected via the reaction graph can be merged. R1 through R8 represent reactions in the network connected by biochemically informed edges, and the orange dashed edge marks a highly correlated reaction pair that cannot merge early on because it lacks a connecting graph edge. The reference dendrogram shows reaction clustering based on reaction correlations across a reference set of cells, resulting in two DDPs (DDP 1 = blue, DDP 2 = green). The perturbed dendrogram shows reaction clustering based on reaction correlations across a perturbed set of cells and the resulting division of DDP 1 into two DDPs. Brackets show the minimum cophenetic correlation between reaction pairs in DDP 1 under the reference and perturbed conditions. **(B)** The maximum Jaccard similarity between each pathway in the top register and the indicated pathway in the bottom register was computed; the mean across all pathways in the top register was then shown. For split controls, cells were randomly divided in half and DDPs were computed independently. **(C)** DDPs were computed at ten minimum-correlation thresholds from 0.05 to 0.90 in enterocytes. For each threshold, coverage was the fraction of all reactions assigned to a DDP, and mean R² was calculated across covered reactions from in-sample OLS models predicting each reaction’s activity from its DDP score. The corresponding KEGG values, obtained using all assigned KEGG pathway scores as predictors, are shown in orange. **(D)** The glycolysis pathway, as defined and laid out in KEGG, is shown. Nodes are metabolites and edges are reactions, with reactions colored by DDP assignment based on enterocytes. A manually chosen subset of metabolites is labeled. **(E)** A superset of panel (D) including all glycolysis reactions and any additional reactions included in the DDPs from panel (D). Yellow nodes indicate metabolites included in panel (D). **(F)** Mean standardized DDP activity across villus regions and proximal-distal domains for the DDPs in panel (E). **(G)** All DDPs overlapping glycolysis, the pentose phosphate pathway (PPP), purine metabolism, or pyrimidine metabolism are shown, with reaction edges colored by DDP assignments as calculated in MS2 cells identified in gastrulating mouse embryos in the accompanying Dias et al. study. **(H)** For each WT MS2 DDP, the weakest link analysis quantified two WT-minus-Folr1 KO metrics: (1) the drop in minimum pairwise reaction correlation and (2) the drop in minimum pairwise cophenetic correlation. DDP 44 is highlighted. **(I)** The relative position of every WT and *Folr1* KO MS2 cell along the cell cycle was computed with Tricycle^89^. For GAR transformylation and AIR carboxylation, reaction activity was smoothed using a centered 200-cell rolling mean and plotted as log2 fold change relative to the mean smoothed activity for that reaction and condition. Background colors indicate cell-cycle phases assigned from Tricycle position.

To test how DDPs differ from canonical metabolic pathways and between cell types, we calculated DDPs for three cell types across two datasets: crypt stem cells and enterocytes from the intestine dataset analyzed above^48^, and the pluripotent epiblast of developing mouse embryos presented in the accompanying manuscript by Dias et al. **(Methods)**. We observed that DDPs were distinct from KEGG pathways **(Figure 3B)**, suggesting that DDPs capture functional metabolic units that are different from preexisting knowledge-based pathways. Furthermore, DDPs from each cell type were different from each other, suggesting that DDPs capture cell type-specific covariation of reactions **(Figure 3B)**. As a negative control, we randomly split each cell type in two and verified that the DDPs computed for each half of the data were highly overlapping **(Figure 3B)**. These results show that DDPs from MeRN’s reaction activity matrix capture unique metabolic subunits depending on the cell type of interest.

We next tested our assumption that DDPs would better capture reaction-level variation than canonical pathways. A key hyperparameter for selecting DDPs is the minimum correlation of any two reactions in the pathway. We calculated the reaction-level variance explained by the set of DDPs computed with varying values of this hyperparameter for each group of cells, along with the variance explained by canonical pathways as represented by KEGG **(Figure 3C, S3A,B, Methods)**. As the minimum correlation increased, the number of reactions included in a DDP decreased, but the reaction-level variance explained increased, generally plateauing around a minimum correlation threshold of 0.7. In comparison, KEGG pathways covered a similarly high proportion of reactions but showed a substantial decline in reaction-level variance explained. These results support the conclusion that our DDPs capture more reaction-level variation than canonical pathways while maintaining high coverage of the metabolic network. Furthermore, they reveal a coverage-variance tradeoff in DDP construction, where DDPs that include a higher fraction of reactions explain less reaction-level variance.

We further leveraged our multimodal alignment over villi in the small intestine to test whether DDPs remain coordinated at the protein abundance level. Specifically, we calculated the average pairwise Spearman correlations of reactions within each pathway based on protein measurements. To strengthen this test, we excluded reaction pairs that shared enzyme-coding genes. DDP-defined groupings showed significantly higher within-group correlations than expected under the empirical null model, and KEGG pathways showed similarly strong within-pathway correlations **(Figure S3C)**. Although DDPs showed a modest increase in within-group correlations over KEGG pathways, limited protein-reaction coverage and villus sampling hampered our ability to robustly compare the two in this setting. Together, these results demonstrate the generalizability of DDPs beyond the transcriptomic level.

A key benefit of the DDP approach is that it groups reactions that are both biochemically linked and appear transcriptionally co-regulated. As such, we hypothesized that MeRN’s DDPs would break down and extend existing KEGG pathways into functional subunits of metabolism. To assess this possibility, we performed a focused analysis of the DDPs from small intestine enterocytes. Given the ubiquitous and essential role of glycolysis across biology and in enterocytes, we focused on how glycolysis-related reactions were reorganized and extended according to MeRN’s DDPs. Visualizing only reactions in KEGG’s glycolysis pathway revealed the pathway was broken into many individual DDPs **(Figure 3D)**, while expanding this visualization to all DDPs with at least one glycolytic reaction revealed a much more extensive network of reactions, including reactions from the TCA cycle, fatty acid metabolism, and amino acid metabolism **(Figure 3E)**. Each one of these DDPs varied uniquely across the villus and proximal-distal axes of the small intestine **(Figure 3F)**, highlighting that reactions within glycolysis may indeed belong to heterogeneous and independently regulated functional units.

Deeper inspection of individual DDPs revealed differential roles for glycolysis in enterocytes throughout the intestine. DDP 91 connected fatty acid metabolism and glycolysis via acetyl-CoA, peaking in the proximal part of the small intestine in alignment with prior descriptions of dietary fatty acid absorption^48,50^. Interestingly, MeRN’s high-resolution view also localized this DDP to the central portion of the villus within this region, suggesting a degree of cell state tuning as enterocytes travel from the crypt to the villus tip. DDP 128 encompassed reactions breaking down dietary carbohydrates into glucose to feed the TCA cycle, peaking in the central small intestine^48,52^ and midway up the villus^59^ (again aligning with previous work).

### DDPs reveal a structural break in *de novo* purine biosynthesis under folate deficiency

In conjunction with the accompanying manuscript by Dias et al., we applied MeRN to mouse gastrulation and early organogenesis, with a focus on folate’s role in germ layer patterning and growth. Our analysis revealed six distinct metabolic states (MS) characterizing the developing embryo, with one state being almost entirely depleted upon genetic disruption of the key transporter Folr1, which severely restricts the embryo’s ability to capture dietary folate^60,61^. We calculated DDPs using the metabolic state of primarily epiblast and pre-gastrulation progenitors (MS2) and analyzed DDP usage across metabolic states and between WT and *Folr1* KO conditions. These analyses revealed major interconnected changes within glycolysis, the pentose phosphate pathway (PPP), and purine metabolism as critical regulators of the neural tube’s expansion and closure defects associated with embryonic folate deficiency (see Dias et al.). Given the direct link between glycolysis, the PPP, and nucleotide metabolism, we visualized the WT DDPs for these pathways, revealing their direct connection via *de novo* nucleotide biosynthesis **(Figure 3G)**.

We further hypothesized that inspecting the stability of the WT DDPs within the *Folr1* KO cells would provide further insight into folate’s role in the developing embryo. For this purpose, we designed a procedure that we call the *weakest link* analysis (named after the British game show; **Methods**). Intuitively, a reaction’s activity must be correlated with the activity of other reactions in its DDP by construction. As such, if we fix DDP definitions based on the WT reaction activities, but compute correlations with KO reaction activities, the lack of correlation identifies reaction pairs that are the least suitably described by WT DDPs and hence may represent control points underlying the reconfiguration of DDPs between the genotypes **(Figure 3A)**. For each DDP, we compute the difference between the minimum within-DDP WT reaction correlation and the minimum within-DDP *Folr1* reaction correlation. We also compute the same metric using the cophenetic correlation to search for correlation drops resulting in large topological changes of the hierarchical clustering used to define DDPs **(Figure 3A)**. These metrics identify WT DDPs containing the reaction pairs that are the weakest links.

Performing the weakest link analysis on our WT embryogenesis DDPs for the *Folr1* KO reaction activities revealed DDP 44 as having a unique combination of large drops in both raw and cophenetic reaction correlations **(Figure 3H)**. This DDP consists of multiple *de novo* purine synthesis reactions, which link the PPP with purine metabolism, starting with glycinamide ribonucleotide (GAR) transformylation, the 10-formyltetrahydrofolate (10-formyl-THF)-dependent conversion of GAR to formylglycinamide ribonucleotide (FGAR) catalyzed by phosphoribosylglycinamide formyltransferase (GART; KEGG R04325), and ending with AIR carboxylation, the conversion of aminoimidazole ribonucleotide (AIR) to carboxyaminoimidazole ribonucleotide (CAIR), as catalyzed by PAICS (KEGG R04209) **(Figure 3G)**. These two reactions showed the largest raw and cophenetic correlation drops in the DDP between WT and KO **(Figure S3D,E)**. Moreover, this drop was associated with a loss of coordination between the reactions over the cell cycle: while both reactions peaked during the DNA synthesis (S) phase in the WT embryo, only GAR transformylation retained a weakened S-phase peak in the KO, with AIR carboxylation losing all coordination with the cell cycle **(Figure 3I)**. Therefore, our weakest link analysis revealed folate deficiency-associated metabolic rewiring between critically affected pathways described in the accompanying manuscript by Dias et al., and can extend preliminary findings to high-resolution detection of critical breakpoints in metabolic topology between control and experimental conditions.

### MeRN-based *metabolic directions* enable metabolic interpretation of cytokine perturbations

Predicting transcriptional changes following genetic, drug, or chemical perturbation is a central goal of machine learning in single-cell genomics^62–65^. However, little attention has been paid to understanding how metabolic state changes in response to these perturbations, or whether metabolic and non-metabolic changes are intrinsically coupled, meaning perturbations that direct similar non-metabolic shifts also drive similar metabolic shifts. We enable this analysis with MeRN by introducing *metabolic directions*, direction vectors from unperturbed to perturbed cells taken within the metabolic latent space. Comparing both direction and magnitude of these vectors enables comparison across perturbations, and the same analyses can be done in the non-metabolic latent space to ultimately compare clustering of perturbations between metabolic and non-metabolic responses. Although these vectors can be calculated on any transcriptional latent space, MeRN offers a unique level of interpretability as the inner product between the metabolic direction vector and each reaction embedding produces a quantifiable metric of how each reaction activity (and consequently DDP) changes upon perturbation **(Figure 1D)**.

We tested MeRN’s ability to interpret immune cell metabolic and non-metabolic responses to cytokine perturbation in the scRNA-seq cytokine dictionary from Cui et al.^66^. The dataset consisted of 23 immune cell types from mouse lymph nodes as they responded to 86 discrete cytokines, with over 300,000 cells profiled by scRNA-seq. Training MeRN and visualizing the metabolic and non-metabolic latent spaces revealed several key differences. While the non-metabolic latent space captured expected cell type differences, the metabolic latent space primarily captured metabolic variation between T cells and myeloid cells **(Figure 4A)**, including increased ether lipid, tryptophan, and glycosaminoglycan metabolism in myeloid cells and increased pyruvate, purine, and propanoate metabolism in T cells **(Figure S4A)**. We then assessed the degree to which cell type versus cytokine treatment dictated metabolic activity by calculating DDPs with cells from the entire dataset and fitting an ordinary least squares model for each DDP **(Methods)**. We found cell type coefficients had higher magnitudes across DDPs than cytokine coefficients, indicating that, at the dataset level, cell type is more impactful in determining metabolic state than cytokine perturbation **(Figure 4B)**. This aligns with the original analysis from Cui et al., which described the cell-type-specific nature of cytokine-induced transcriptional changes.

**Figure 4.**
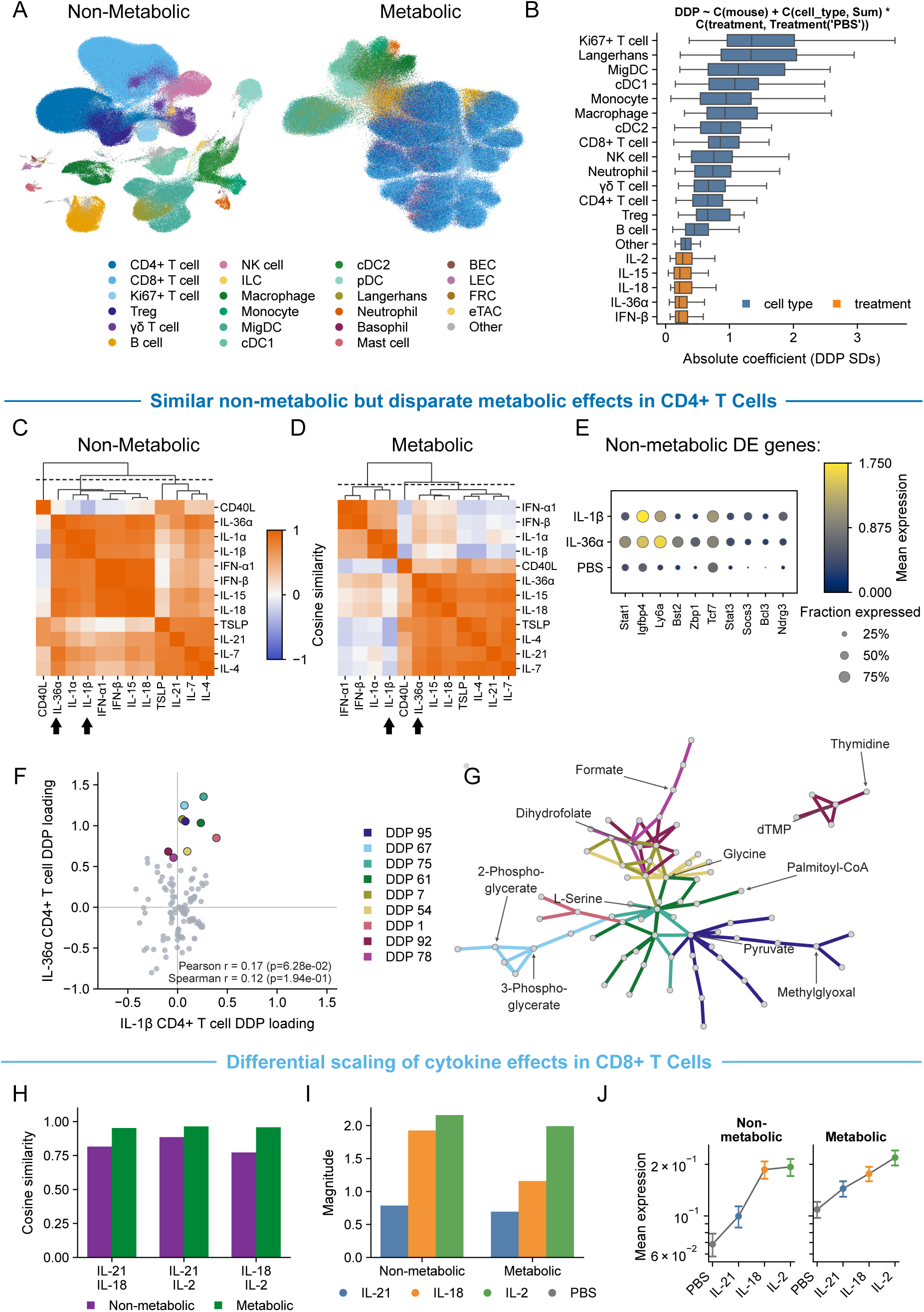
MeRN disentangles metabolic and non-metabolic responses to cytokine perturbation. **(A)** UMAP representations of non-metabolic and metabolic MeRN latent spaces, colored by cell type. **(B)** DDP activity was averaged within mouse-cell type-treatment groups and groups with 25 or fewer cells were excluded. For each DDP, activity was standardized over the remaining groups, and an ordinary least-squares model was fit including mouse, sum-coded cell type, PBS-referenced treatment, and the cell type-by-treatment interaction. For visualization, effects with at least 10 FDR-significant DDP associations were eligible for display, and the top effects were selected by mean absolute coefficient and ordered by median absolute coefficient. Blue represents a cell-type effect; orange represents a cytokine-treatment effect. **(C)** Cosine similarity between non-metabolic direction vectors for CD4+ T cells. Cytokines were ordered by average-linkage hierarchical clustering of the similarity matrix. The dashed line marks the dendrogram height that produces three clusters. **(D)** Same as panel (C), except for metabolic direction vectors. **(E)** For CD4+ T cells treated with IL-1β, IL-36α, or PBS, color indicates mean expression, and dot size indicates the fraction of cells expressing each selected non-metabolic gene. Displayed genes were manually selected from the top 20 significantly upregulated non-metabolic genes for IL-36α (left 5 genes) and IL-1β (right 5 genes). **(F)** DDP loadings were computed by multiplying the metabolic graph embedding by the IL-1β or IL-36α CD4+ T-cell direction vector and then averaging reaction loadings within each DDP (Figure 1D). DDPs with IL-36α loading greater than 0.6 are highlighted. **(G)** Pathway network plot for the DDPs highlighted in panel (F). Reaction edges are colored by DDP membership. **(H)** Cosine similarities of metabolic and non-metabolic CD8+ T-cell direction vectors for the three pairwise comparisons among IL-2, IL-18, and IL-21. **(I)** Magnitude of non-metabolic and metabolic direction vectors for IL-2, IL-18, and IL-21, calculated as the L2 norm of each vector. **(J)** Mean log-normalized expression of the top 100 significantly upregulated metabolic and non-metabolic genes identified from the combined IL-2, IL-18, and IL-21 versus PBS CD8+ T-cell contrast; error bars show the standard error across genes.

Our results motivated us to develop methods for extracting cell type-specific metabolic directions with MeRN. Specifically, we decided to explore cell type-specific metabolic responses to cytokine stimulation in CD4+ and CD8+ T cells by calculating metabolic directions for each cell type from unperturbed (PBS) cells to all profiled cytokines. We used the cosine distance between direction vectors for biological replicates to select perturbations with consistent effects. While low cosine distance was associated with larger direction magnitude, as expected, perturbations with a range of magnitudes showed low cosine distance compared to the negative control (PBS), and we retained 12 and 15 unique cytokines in CD4+ and CD8+ T cells, respectively, for downstream analysis **(Figure S4B,C)**.

To compare metabolic and non-metabolic effects across cytokines in each cell type, we calculated pairwise cosine similarities among the metabolic and among the non-metabolic directions. CD4+ T cells showed distinct clusters of cytokines based on their non-metabolic and metabolic direction vectors, primarily resolving into three groups for each test **(Figure 4C,D)**. However, these three binned subsets of responses were not the same for metabolic and non-metabolic responses, indicating that metabolic and non-metabolic effects of cytokines are not necessarily paired for CD4+ T cells. For example, IL-36α and IL-1β had similar non-metabolic direction vectors but disparate metabolic direction vectors **(Figure S4D)**. Plotting selected significantly upregulated non-metabolic genes for IL-1β and IL-36α confirmed the similarity of responses with genes such as *Stat1* and *Tcf7* being increased under both conditions **(Figure 4E)**. In contrast, the changes in DDP activity induced by IL-1β and IL-36α showed low correlation, with many DDPs increasing much more after IL-36α treatment **(Figure 4F, Methods)**. Plotting the DDPs that were strongly induced by IL-36α but only weakly induced by IL-1β suggested a strong proliferative response to IL-36α in CD4+ T cells, with folate, serine, and pyrimidine metabolism being upregulated **(Figure 4G)**. In all, these results demonstrate the value of characterizing metabolic and non-metabolic responses separately, as we show that IL-1β and IL-36α induce shared non-metabolic responses but differential metabolic responses in CD4+ T cells. Furthermore, MeRN enables interpretation of metabolic responses as changes in DDP activity, generating testable hypotheses for the differences between CD4+ T cell metabolic responses to IL-1β and IL-36α.

We then explored cytokine-induced metabolic and non-metabolic effects in CD8+ T cells. Similarly to CD4+ T cells, cytokine-induced changes formed distinct clusters for both non-metabolic and metabolic directions **(Figure S4E,F)**. Rather than further exploring differential clustering, we focused on another aspect of cellular response: magnitude. In this context, the magnitude of the direction vector describes the strength of the metabolic or non-metabolic transcriptional response. Within both the metabolic and non-metabolic latent spaces, the direction vectors induced by IL-21, IL-18, and IL-2 had high cosine similarity but differing magnitudes across cytokines **(Figure 4H,I)**. Interestingly, we observed a stepwise increase in the magnitude of the non-metabolic direction vectors, with a twofold increase from IL-21 to IL-18 followed by a much smaller increase from IL-18 to IL-2. In contrast, the metabolic magnitudes increased more gradually across cytokines. Both trends could be confirmed by plotting the average expression of the top 100 non-metabolic and metabolic differentially expressed genes **(Figure 4J)**, highlighting that our analysis captures differential response scaling of metabolic and non-metabolic cytokine effects. Together, these results establish the ability of metabolic directions to reveal the type and magnitude of metabolic response to CD4+ and CD8+ T cell cytokine perturbation. This capability could enable targeted cytokine selection and engineering by distinguishing cytokines with similar immune-state effects but disparate metabolic programs.

### MeRN reveals pan-cancer metabolic drivers of T cell function

Having established MeRN’s ability to describe metabolism across diverse biological contexts and given the importance of metabolism for immune cell function in cancer^67^, we sought to leverage MeRN’s scalability to discover translationally relevant metabolic drivers of T cell function and dysfunction in cancer. We applied MeRN to a human atlas of tumor-infiltrating T cells curated by Zheng et al.^68^ and encompassing 397,810 human T cells from 21 cancer types.

Given the complexity of the dataset, we opted for a linear mixed model (LMM) to model DDP activity across T cells, treating T cell type (as defined by Zheng et al.), tissue, and cancer type as fixed effects, and patient as a random effect **(Methods)**. We fit an LMM for 119 DDPs using post-filter pseudobulks from 166 of the 260 patients, 41 cell types, and four tissues, including blood, normal tissue, tumor, and lymph node. We clustered DDPs based on their coefficients across a CD8+ T cell path from naive to effector to exhausted as determined in Zheng et al. **(Figure 5A)**. This revealed five distinct groups of DDPs representing different patterns across the T cell exhaustion trajectory. One cluster, which we termed Exhausted Decrease, had coefficients that decreased throughout the exhaustion trajectory, suggesting these units of metabolism are associated with the naive-like CD8+ T cell state. Two other clusters, namely Exhausted Increase and Exhausted Mild Increase, showed the opposite trend with coefficients increasing throughout exhaustion at different levels of intensity. We further identified a DDP cluster, Effector Associated, which increased specifically in effector and memory CD8+ T cells. The remaining DDPs were not clearly associated with the effector-to-exhaustion trajectory, which we termed Not Associated.

**Figure 5.**
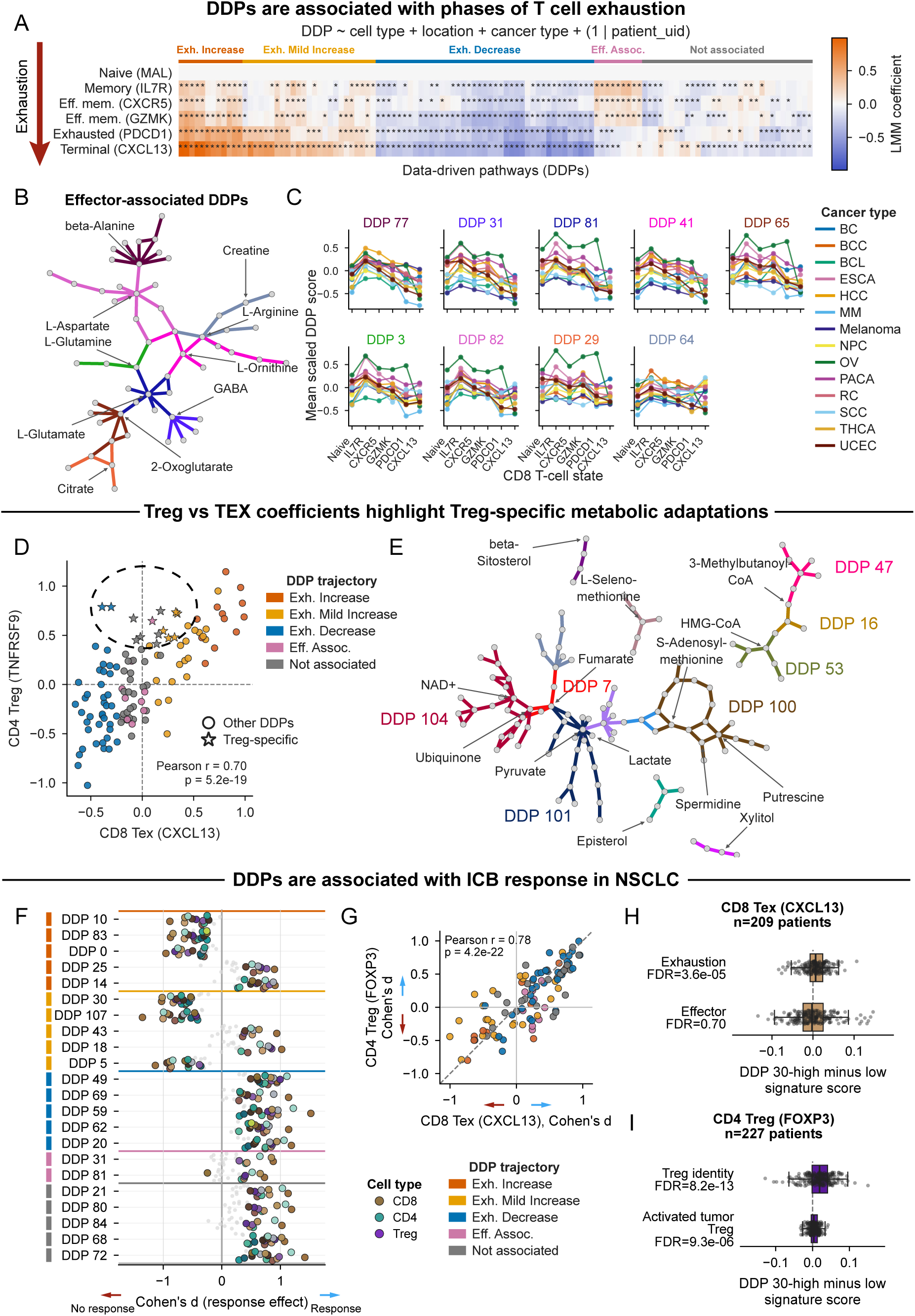
MeRN reveals metabolic drivers of pan-cancer human T cell function. **(A)** DDP activities were pseudobulked by patient, cancer type, tissue location, and cell type; groups with 25 or fewer cells were excluded. For each DDP, an LMM included fixed effects for cell type, tissue, and cancer type and a random intercept for patient. Cell-type coefficients are displayed. Cell types were ordered along a trajectory of CD8+ T cell exhaustion from Zheng et al.^68^, and asterisks mark Benjamini-Hochberg-adjusted p < 0.05 within the cell-type effect family. DDPs (columns) were clustered by hierarchical clustering. **(B)** Pathway network plot of effector-associated DDPs, with reactions colored by DDP membership. **(C)** Mean scaled pseudobulked DDP activity across the six-state CD8+ T cell exhaustion trajectory for each cancer type. **(D)** CD8 Tex (CXCL13) versus CD4 Treg (TNFRSF9) LMM coefficients. Stars identify Treg-specific DDPs that pass three one-sided Wald tests, each at Benjamini-Hochberg FDR < 0.05: the Treg coefficient is greater than 0.35, the Tex coefficient is less than 0.45, and the Treg coefficient exceeds the Tex coefficient. **(E)** Pathway network of the Treg-specific DDPs identified in panel (D). Reactions are colored by DDP membership. **(F)** Patient-level DDP activity was compared between responders and non-responders within each cell type. Patient-cell type groups with 25 or fewer cells were excluded, and each analyzed cell type required at least 25 responders and 25 non-responders. DDPs significant at Benjamini-Hochberg FDR at or below 0.05 in at least six cell types were retained; panel (F) shows up to five DDPs per trajectory cluster with the largest significant absolute Cohen’s d. Each point represents Cohen’s d for a given DDP (row) and cell type (color). Colored points indicate FDR-significant effects; smaller gray points are not significant. CD4, CD8, and Treg colors are summarized with the full legend in Figure S5. **(G)** CD8 Tex (CXCL13) versus CD4 Treg (FOXP3) Cohen’s d values for all shared DDPs, colored by the DDP trajectory clusters from panel (A). The identity line and Pearson correlation summarize effect-size alignment. **(H)** The CD8 T-cell dysfunction/exhaustion and cytotoxic-effector gene sets from Gavish et al.^91^ were used to score CD8 Tex (CXCL13) cells from each patient. For each patient with more than 25 cells, the mean score in the DDP 30-low quartile was subtracted from the mean in the DDP 30-high quartile. P values were calculated by a one-sided Wilcoxon signed-rank test across patients and corrected jointly across the four signatures with the Benjamini-Hochberg procedure. **(I)** Same as panel (H), but using the Treg-identity gene set from Gavish et al.^91^ and an activated-tumor-Treg gene set derived from Alvisi et al.^90^.

We further investigated the Effector Associated DDP cluster corresponding to activated and effector CD8+ T cell subsets. These DDPs consisted of reactions connecting glutamine to the TCA cycle, along with arginine-centered nitrogen metabolism involving creatine and ornithine **(Figure 5B)**. Notably, arginine and creatine metabolism have been shown to be essential for T cell activation and function^69,70^, supporting the finding of these DDPs as effector-related. Furthermore, glutaminolysis is necessary for T cell activation and function, and is downregulated during exhaustion^71,72^. Further examination of the reaction between glutamine and glutamate revealed interesting nuances. The reaction converting glutamine to glutamate (the direction of glutaminolysis) is canonically driven by glutaminase (GLS), while the reverse direction is catalyzed by glutamate-ammonia ligase (GLUL). Although MeRN does not distinguish between these directions, examining the weights of the reaction-to-gene layer reveals that the learned reaction activity is largely driven by GLUL rather than GLS, suggesting a potentially distinct mechanism from canonical glutaminolysis **(Figure S5A)**. Finally, plotting the activity of these DDPs over the exhaustion trajectory separately for each cancer type revealed the consistency of this upregulation specifically in effector and memory CD8+ T cells, before decreasing in exhaustion **(Figure 5C).** Together, these results demonstrate the ability of MeRN-based DDPs to characterize pan-cancer metabolic support for CD8+ T cell activation and effector function.

We next turned to identifying differences in metabolic function between two T cell subtypes characterized as tumor-enriched and potentially tumor-reactive in Zheng et al.: TNFRSF9+ regulatory T cells (Tregs) and terminally exhausted CD8+ T cells. Tregs metabolically adapt to the tumor microenvironment (TME) in ways CD8+ T cells cannot, often resulting in immunosuppressive TMEs and poor outcomes for patients^73^. DDP coefficients showed a strong correlation between exhausted CD8+ T cells and tumor-infiltrating Tregs, indicating a shared metabolic program in which both rely on many of the same pathways for activation and survival in the TME. Specifically, the DDPs that increased throughout the CD8+ T cell exhaustion trajectory were also positively associated with Tregs, and the DDPs that decreased were negatively associated **(Figure 5D)**. Notably, we identified 14 DDPs with significantly larger coefficients in Tregs than exhausted T cells (**Figure 5D**, **Methods**). These trends suggest largely shared metabolic adaptations to the TME between Tregs and CD8+ exhausted T cells, with a subset of additional Treg-specific metabolic adaptations.

We then investigated the metabolic programs associated with Treg-specific metabolic adaptation to the TME. These DDPs included reactions in oxidative phosphorylation (DDP 104), TCA cycle (DDPs 101, 7), polyamine metabolism (DDP 100), and fatty acid metabolism (DDPs 47, 16, 53) (**Figure 5E**). DDP 101 specifically highlighted a link between lactate, pyruvate, and fumarate. Tumor Tregs’ ability to absorb lactate via MCT1 for use in the TCA cycle has been previously characterized^74^, and our data shows a strong correlation between DDP 101 activity, which contains the lactate-pyruvate link, and MCT1 expression, with the highest DDP 101 cells also having high *FOXP3* expression **(Figure S5B)**. Together, our results provide a coherent description of pan-cancer CD8+ T cell effector function and exhaustion, along with the identification of novel and literature-supported Treg-specific metabolic adaptations that could form the basis for precision therapeutics targeting Tregs in the TME.

### T cell DDP activity is associated with immunotherapy outcome in NSCLC

We observed heterogeneity in the metabolic states of CD8+ exhausted T cells across patients and cancer types **(Figure S5C)**, prompting us to investigate whether the metabolic states of T cell subtypes are indicative of patient outcome in cancer immunotherapy. We leveraged existing scRNA-seq data from Liu et al.^75^, who profiled over 1 million immune cells from 234 anti-PD-1-treated non-small cell lung cancer (NSCLC) tumors. We trained MeRN on this dataset, allowing us to assess the metabolic states of T cells in responders vs non-responders to anti-PD-1 therapy and associate these metabolic states with the DDPs defined above. The majority of our previously defined T cell atlas DDPs showed high within-DDP correlation across NSCLC T cells, validating the reproducibility of our T cell DDPs **(Figure S5D)**. We therefore focused on these highly correlated DDPs for the analysis of this dataset.

Liu et al. demonstrated that immune cell composition, particularly a high frequency of Tregs, was highly indicative of anti-PD-1 non-response. We reasoned that individual cell states should also differ between responders and non-responders. To assess this, we fit logistic regression models to predict patient outcome from the average DDP values for each T cell subtype of each patient. As a baseline, we also predicted patient outcomes by the average non-metabolic embedding of the same cell subtypes. Both metabolic and non-metabolic cell type features showed high AUCs across many cell types, including exhausted CD8+ T cells and Tregs, suggesting that the states of specific immune cell types, and not only their composition, are highly predictive of patient response to immunotherapy in this dataset **(Figure S5E)**.

Given the predictive power of metabolic state as defined by average DDP activity across many T cell subtypes, we investigated whether specific DDPs could be associated with patient outcome. Within each T cell subtype, we identified many DDPs that had a significant association with response, with pro- or anti-response effects being largely consistent across T cell subtypes (**Figure 5F, S5F)**. DDPs that decreased throughout exhaustion or were associated with effector function were generally increased in the T cells of responders. In contrast, DDPs positively associated with exhaustion were generally decreased in the T cells of responders (**Figure 5F, S5F)**. Strikingly, we noticed a strong positive correlation between the differential usage of DDPs between responders and non-responders within CD8+ exhausted T cells and Tregs, indicating that DDPs were associated with response or non-response consistently across the two cell types **(Figure 5G)**. We found this surprising given the opposing roles of Tregs and CD8+ T cells in the TME and hypothesized that this could be explained by non-responder-associated DDPs inducing exhausted CD8+ T cells with a stronger exhausted phenotype, while simultaneously inducing more suppressive Tregs. Comparing DDP 30-low and DDP 30-high CD8+ exhausted T cells and Tregs confirmed that DDP 30-high CD8+ TEX cells were indeed more exhausted without showing significant differences in effector-linked gene expression. Simultaneously, DDP 30-high Tregs had higher expression of Treg identity markers and genes associated with activation of Tregs in the tumor **(Figure 5H,I)**. These results support a fundamental role for metabolism in determining immunotherapy outcomes^76–78^ and suggest that metabolic pathways have a synergistic effect in supporting effector function in CD8+ T cells and impairing suppressive Treg function.

## Discussion

Here, we introduce MeRN, a graph-guided VAE that leverages prior knowledge of the complex metabolic network to learn interpretable single-cell metabolic states from scRNA-seq data. Our approach bridges reaction- and pathway-level frameworks for inferring metabolic state from transcriptomic data, enabling the construction of metabolic DDPs as an unbiased, data-driven approach to understanding cellular metabolism. We demonstrate MeRN’s scalability, robustness, and ability to enable novel insights across a variety of biological settings.

First, we benchmarked MeRN against pathway- and reaction-level analyses in the small intestine. By curating data on the same biological system across multiple datasets, we enabled comprehensive characterization of cross-modality measurements of enterocyte metabolism. MeRN rediscovered known trends in enterocyte metabolism and accurately predicted metabolic activity at the pathway and reaction levels. Furthermore, MeRN correlated most strongly with protein measurements among all methods, suggesting its ability to leverage prior knowledge of the metabolic topology to learn reaction activities closer to protein expression, and as a result, true metabolic activity.

Second, we identified the need for data-driven partitioning of metabolism into functional subunits, in lieu of predefined pathways, and leveraged MeRN’s complete metabolic representation to construct cell state-specific DDPs. Predefined pathways assume a static partition of metabolic functions; however, the organization of the metabolic network into functional subunits can change among cell types, disease states, and experimental conditions. We showed our learned DDPs vary across cell types and datasets and differ significantly from predefined KEGG pathways. Investigating glycolysis-related DDPs in enterocytes highlighted the ability of DDPs to extend existing pathways and capture physiologically relevant subunits of metabolism associated with nutrient absorption and intestinal function. We then established the weakest link analysis as a novel method to computationally generate mechanistic predictions of metabolic rewiring based on DDPs. Applying this analysis to wild-type and *Folr1* mutant data during embryogenesis revealed rewiring of *de novo* purine synthesis associated with depletion of a specific metabolic cell state linked to embryonic neural tube defects. More generally, with the weakest link analysis, we move beyond differential gene expression, reaction activity, or pathway activity, enabling analysis of metabolic reprogramming as structural reorganization of pathways within the metabolic network.

Third, we introduced MeRN-based metabolic directions for interpretable analysis of metabolic perturbation effects and applied this technique to understand cytokine-induced changes in different immune cells. We found IL-36α and IL-1β induced shared effector responses but highly divergent metabolic responses in CD4+ T cells. The link between MeRN’s metabolic latent space and graph embedding enabled interpretation of these metabolic responses as changes to DDP activity, generating the testable mechanistic hypothesis that proliferation-associated pathways may explain these divergent metabolic responses. Furthermore, we demonstrated differential scaling of metabolic and non-metabolic transcriptional responses in CD8+ T cells. Overall, metabolic directions provide a powerful toolkit for interrogating the often-overlooked metabolic component of perturbation responses. More broadly, this framework could prioritize cytokines, ligands, and genetic perturbations that induce a desired non-metabolic cell-state effect with favorable biosynthetic or energetic programs.

Finally, we applied MeRN to a pan-cancer human T cell atlas, revealing metabolic DDPs associated with different stages of CD8+ T cell exhaustion. Our results showed that usage of these DDPs was largely shared between tumor-associated Tregs and CD8+ exhausted T cells, with a subset of additional Treg-specific metabolic adaptations. These DDPs may be associated with the ability of Tregs to maintain their suppressive functionality in the TME, whereas CD8+ T cells progressively lose their cytotoxic functionality. Therefore, this Treg-specific subset represents a promising avenue for specifically targeting Tregs in the TME. Furthermore, our identified DDPs were reproducible in an independent dataset of immune cells from NSCLC tumors following anti-PD-1 therapy. We showed that DDP activity was associated with clinical response to anti-PD-1 therapy consistently across T cell subtypes, often with the expected direction of effect given their association with T cell exhaustion from the previous dataset. By identifying nuanced metabolic programs that support both suppressive Tregs and exhausted CD8+ T cells, we uncovered promising therapeutic targets. This contrasts with traditional immune checkpoint inhibitors, where anti-PD-1 can simultaneously reinvigorate PD-1+ effector T cells and enhance the suppressiveness of PD-1+ Tregs in NSCLC^79^.

This work provides a foundation for the future development of methods that integrate prior biological knowledge (in the form of complex graphs) with high-resolution measurements of cells to learn interpretable cell states. We extend beyond associative networks such as gene-gene graphs and leverage a biochemical prior in the form of the metabolic network. Within scRNA-seq, the metabolic graph could be extended to include transporters or other regulatory layers such as epigenetic state, or replaced with other networks representing gene-gene relationships, such as those from gene regulatory networks^44,63,80,81^ or cell-cell communication^82–85^. Furthermore, extending MeRN to input other modalities such as spatial and epigenetic state offers promise for understanding the bidirectional regulatory mechanisms that govern cellular function and state.

The introduction of MeRN-based DDPs is a step towards decomposing the complex metabolic network based on observed variation. While existing pathway definitions are highly useful, they overlook context-dependent heterogeneity common in disease settings. Existing methods for defining data-driven metabolic modules, such as GAM-clustering^57,58^, identify transcriptionally coordinated subnetworks, but only produce a limited number of modules covering a small portion of the metabolic network in reported applications^57,58^. In contrast, MeRN-based DDPs provide a high-coverage decomposition of the metabolic network, both constrained by the biochemical connectivity of reactions and informed by cellular context. Furthermore, they allow for complex network-based analyses such as investigating rewiring under perturbed conditions, as demonstrated by identifying a structural break in *de novo* purine synthesis under folate-depleted conditions in the developing embryo. Extending DDPs to include other modalities or molecular interaction networks is a promising future direction for elucidating context-dependent biological regulation.

## Limitations of the study

MeRN’s metabolic topology graph was constructed from organism-specific KGML representations of the KEGG global metabolic pathway map (KEGG Release 113.0-era snapshot, generated January 15, 2025). This manually curated and interpretable representation sacrifices stoichiometry, biochemical reaction directionality, and subcellular metabolite compartmentalization. As a result, the graph contains a representative subset of cellular metabolism and generally overlooks metabolite transporters, although they are informative. Additionally, reaction activity is not indicative of reaction directionality. MeRN relies on transcriptional measurements of enzyme-coding genes, thereby overlooking post-transcriptional and post-translational regulation, which may be especially important for short time-scale adaptations. We enforce a strict cutoff between metabolic and non-metabolic genes, overlooking potentially unannotated metabolic enzymes and the non-metabolic roles of moonlighting enzymes.

## Resource Availability

MeRN is implemented as a Python package available at https://github.com/wagnerlab-berkeley/mern, along with a comprehensive tutorial. Scripts and data for reproducing all figures in this manuscript will be made available upon publication.

## Acknowledgments

We thank members of the Wagner and Ioannidis labs, especially Charles Chien and Ayesha Bajwa, for useful feedback throughout model development and the writing process. We thank Yun Song and Ellen Robey for helpful discussions and feedback on the work. DPL, AC, and AW were supported by a research grant from the Shurl and Kay Curci Foundation. NMI is a Biohub San Francisco Investigator. ZDS is supported by the National Institutes of Health (NIH) Director’s New Innovator Award (DP2HD108774), the Mathers Foundation, the Chen Innovation Award, and the Max Planck Society.

## Author Contributions

DPL, NMI, and AW conceptualized and designed MeRN. DPL, ND, AC, ZDS, NMI, and AW conceptualized the analytical methods, applications, and validation of MeRN. DPL and AC implemented the MeRN software. DPL, ND, AC, and YK collected, analyzed, and interpreted the data. NMI, ZDS, and AW supervised the work. DPL, ND, ZDS, NMI, and AW wrote the manuscript with input from all authors. All authors reviewed and approved the final manuscript.

## Declaration of generative AI and AI-assisted technologies in the writing process

The manuscript was written by the authors, and AI was used to improve the grammar and clarity of the writing. For figure legends and methods, agentic AI with access to the codebase was used to verify that the textual description fully aligns with the codebase. All authors reviewed the final manuscript and take full responsibility for its contents.

## Methods

### Datasets and Preprocessing

#### Shared Preprocessing

Except where noted for the T cell atlas, all datasets were preprocessed using the same workflow. First, genes detected in fewer than 100 cells were removed. Highly variable genes were then identified separately within the metabolic and non-metabolic gene sets using the scanpy implementation of the seurat_v3 method. The top 5,000 most variable non-metabolic and 800 most variable metabolic genes (defined below) were retained for model training. For the T cell atlas, 3,000 non-metabolic genes and all 748 metabolic genes remaining after gene filtering were retained because relatively few genes remained after atlas preprocessing. We calculated highly variable genes separately for metabolic and non-metabolic genes to increase the number of metabolic genes available to the model. Further dataset-specific details are provided below.

### Small intestine

After preprocessing, the dataset contained 19,847 cells. No batch covariate was used for model training. Cell-type labels and domain assignments were obtained from the original dataset^48^.

### Embryogenesis

After preprocessing and subsetting to core embryonic tissues by excluding extraembryonic ectoderm, extraembryonic endoderm, and blood, the wild-type (WT) and *Folr1* KO datasets contained 62,433 and 31,771 cells, respectively. Separate MeRN models were trained for WT and Folr1 cells with no batch covariate. Cell types were assigned as described in the accompanying manuscript by Dias et al.

### Cytokine Dictionary

After preprocessing, the dataset contained 386,703 cells. The channel annotation was included as a batch covariate during model training, and cell-type annotations were obtained from the original dataset.^66^

### T cell atlas

We downloaded dataset-specific SingleCellExperiment objects curated and released by Zheng et al.^68^ For datasets with raw counts, we extracted counts and metadata from the objects provided with the paper. To create a merged AnnData object, we constructed a global set of genes shared across all datasets based on the display.name identifier. After preprocessing, the dataset contained 373,380 cells and 9,475 genes. The dataset annotation was included as a batch covariate during model training, and cell-type annotations were obtained from the original study.

### NSCLC Immune Cell Atlas

After preprocessing, 1,254,749 cells were used for model training. No batch covariate was used during model training, and cell-type annotations were obtained from the original study^75^.

### Metabolic Guidance Graph

#### KGML-Based Metabolic Guidance Graph Construction

We leveraged KEGG to construct our undirected, unweighted metabolic guidance graph *G* = (ℛ, ℰ), where reactions (ℛ) are nodes and edges (ℰ) are shared metabolites between reactions. For each species, we downloaded the global pathway map (01100), which contains 1,270 and 1,251 reaction elements (1,264 and 1,245 unique reaction-entry names) in the archived human and mouse metabolic maps, respectively. Each KEGG Markup Language (KGML) reaction entry contains one or more space-separated KEGG reaction identifiers, and each unique reaction-entry name defines one graph node. The KGML also includes the substrates and products displayed for each reaction entry. Because we used only these displayed metabolites to construct graph edges, omitted metabolites—including many occurrences of common cofactors such as water and ATP—did not contribute to graph connectivity. We also manually added five oxidative phosphorylation reactions omitted from the global pathway map (R11945, R13223, R13224, R02161, and R00081).

We constructed an undirected metabolic topology graph *G* by adding edges (*i*, *j*) ∈ ℰ between distinct reaction entries that share any displayed metabolites (either products or substrates). We treated substrates and products for each reaction as an unordered set because KGML reaction entries often contain reversible reactions or combine forward and reverse reactions. KGML entries with identical reaction-entry names were represented by the same graph node. Finally, we removed isolated reactions that share no neighbors in the graph. Our final human metabolic graph includes 1,225 reaction nodes and 2,802 undirected edges. Our final mouse metabolic graph includes 1,213 reaction nodes and 2,791 undirected edges. Both are available through our MeRN Python package. These choices were intended to construct a metabolic topology graph that captures pathway-level relationships rather than model biochemical flux.

### Reaction-to-Gene Mapping

To map each reaction node to genes encoding enzymes that catalyze its constituent reaction or reactions, we first pooled the KEGG enzyme entries linked to all KEGG reaction identifiers in that node. We then linked these enzymes to species-specific KEGG gene identifiers and converted the identifiers to gene symbols using the Python mygene package separately for human and mouse. Before model fitting, we deduplicated each gene list and restricted it to genes present in the input transcriptomic dataset. This process produced a list of available enzyme-coding genes for each reaction node. Because we do not distinguish between compartmental localization of reactions, our reaction-to-enzyme mapping includes ectoenzymes such as NT5E and ENTPD1.

### MeRN Model and Training Model Description

MeRN is a graph-guided generative model that uses the metabolic topology graph (constructed as described above) as an inductive bias to learn interpretable single-cell metabolic states. It achieves this by learning separate metabolic and non-metabolic latent spaces, which reconstruct metabolic (enzyme-coding) and non-metabolic genes, respectively. The metabolic topology graph serves to make the metabolic latent space interpretable with respect to reactions in the metabolic graph and enable estimation of reaction activity.

Our use of a guidance graph was inspired by GLUE^44^, a method for integrating single-cell multiomic datasets. GLUE leverages a guidance graph linking features across input modalities based on prior knowledge of the gene regulatory network, enabling joint embedding of single cells from distinct modalities. Below, we adapt the graph likelihood (Equation 3), the use of a graph convolutional network (GCN) encoder (Equation 11), and the decoder linking cell and feature embeddings (Equation 5). Unlike GLUE, MeRN’s guidance graph represents a biochemical inductive bias over reactions rather than input molecular features. Furthermore, we introduce separate metabolic and non-metabolic latent spaces that reconstruct disjoint gene sets, a nonnegative reaction activity layer in the decoder for mechanistic interpretability, and a masked nonnegative linear layer based on reaction-enzyme annotations.

We use *x* ∈ ℝ^|S|^ to denote the gene-count vector for a cell measured by scRNA-seq, where S is the set of genes. We also introduce the metabolic guidance graph *G* = (ℛ, ℰ), which represents prior knowledge about the metabolic topology. ℛ is the set of reactions and ℰ is the set of edges. Let *z* denote the shared dimensionality of the metabolic and reaction embeddings, and *u* denote the dimensionality of the non-metabolic latent space, where both *z* and *u* are model hyperparameters.

Each reaction *i* ∈ ℛ is assigned a latent variable *v_i_* ∈ ℝ*^z^*, and we collect these embeddings as the rows of *V* ∈ ℝ^|ℛ|×*z*^. We then define the conditional probability of the metabolic guidance graph given *V*.

We model the observed scRNA-seq data for each cell as generated from a metabolic latent space *m* ∈ ℝ*^z^*, a non-metabolic latent space *b* ∈ ℝ*^u^*, and the metabolic reaction latent space *V*. The combined model likelihood can be defined as follows:

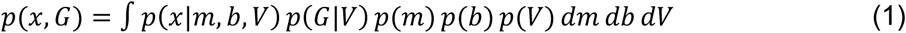

where *p*(*x*|*m*, *b*, *V*) and *p*(*G*|*V*) specify the observation models for cellular transcriptomic profiles and the metabolic graph, respectively. We place standard normal priors on the metabolic latent space (*m*), non-metabolic latent space (*b*), and reaction (graph) latent space (*V*).

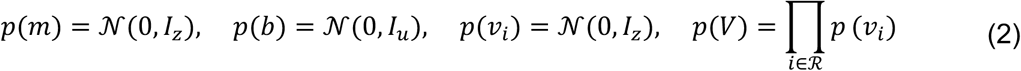

We reconstruct the metabolic graph by parameterizing a Bernoulli distribution with negative sampling based on pairwise similarities between reaction embeddings:

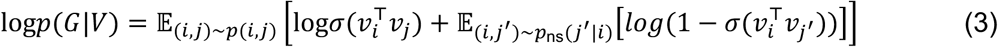

Here, *σ*(*x*) represents the sigmoid function and *p*_ns_ is the sampling distribution used to select nodes not connected to *i*. Importantly, our graph decoder has no learnable parameters. We define the scRNA-seq data likelihood *p*(*x*|*m*, *b*, *V*) as follows:

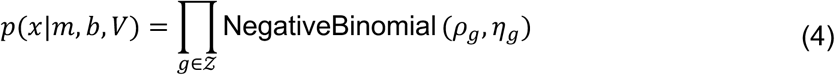

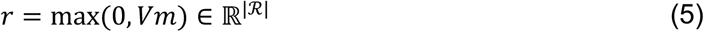

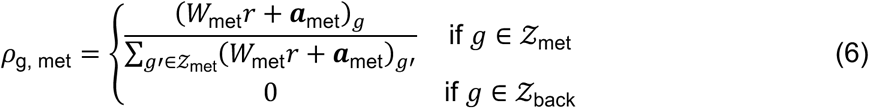

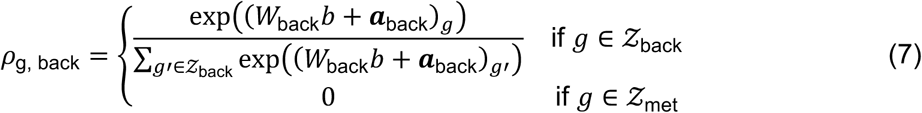

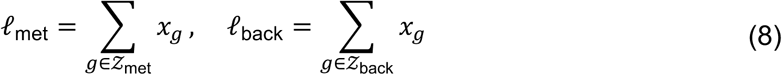

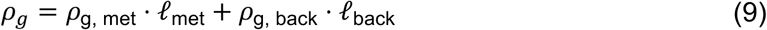

Here, *ρ*, *η* ∈ ℝ^|z|^ are the mean and dispersion parameters of the Negative Binomial distribution. S_met_ refers to the subset of genes encoding metabolic enzymes, and S_back_ refers to the subset of genes not encoding enzymes. *W*_met_ and ***a***_met_ are the weight matrix and gene-specific bias vector of the reaction-to-gene layer. Its weights are masked according to the reaction–gene annotations, and both the weights and bias are clamped to be nonnegative during the forward pass. Thus, through the reaction weights, each reaction can contribute directly only to its associated enzyme-coding genes (reaction–gene curation is described above). *W*_back_ and ***a***_back_ are the corresponding weight matrix and bias vector for the background decoder, whose parameters are not constrained to be nonnegative. This probabilistic model defines the count of each gene in a given cell as generated from metabolic and non-metabolic components, with metabolic and non-metabolic gene library sizes treated separately.

Because the true posterior is intractable, we approximate it with a mean-field variational posterior:

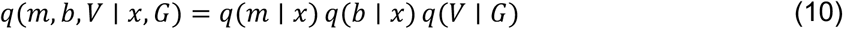

A GCN dependent on *G* predicts the mean and diagonal covariance of the variational posterior for each reaction:

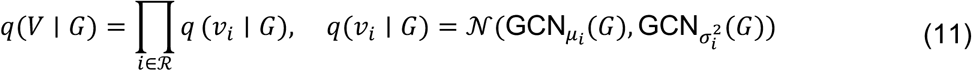

Separate multilayer perceptron (MLP) neural networks specific to the metabolic and non-metabolic latent spaces map gene expression *x* to the mean and diagonal covariance of metabolic and non-metabolic cell embeddings:

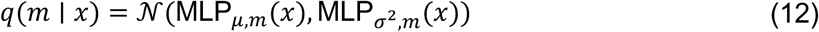

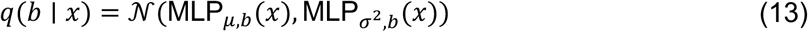

Applying Jensen’s inequality to the model described above, we can derive the evidence lower bound (ELBO):

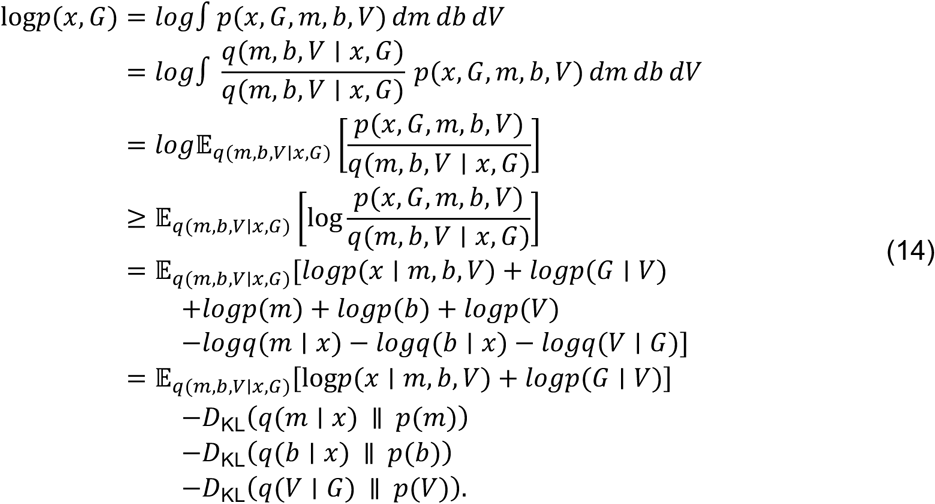

Across a dataset of *n* cells, we denote the collections of metabolic and non-metabolic cell embeddings by *M* ∈ ℝ*^n^*^×*z*^ and *B* ∈ ℝ*^n^*^×*u*^, respectively. The lowercase *m* and *b* above denote the embeddings for an individual cell. The matrix of decoded reaction activities across these cells is *R* = max(0, *MV*^T^) ∈ ℝ*^n^*^×|ℛ|^, where the maximum is applied elementwise.

### Model Training

For model optimization, we minimize the negative empirical counterparts of the data ELBO loss (ℒ_data_) and graph ELBO loss (ℒ_graph_). We normalize the reconstruction losses by the number of genes (|S|) and the KL divergences by their respective latent dimensionalities (*z* and *u*). The graph reconstruction term is averaged over sampled edges, and the reaction-embedding KL term is averaged over reaction nodes. We also introduce *β*-weights to scale the KL divergence terms:

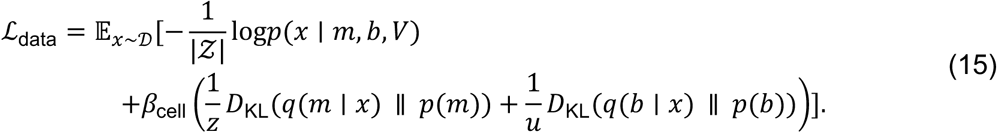

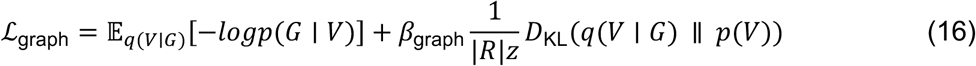

Finally, the total empirical loss used for stochastic optimization is a weighted linear combination of these two components, governed by hyperparameters (*λ*):

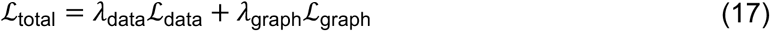

All models in this manuscript were trained with latent dimensions *z* = 25 and *u* = 15, KL divergence weights *β*_cell_ = 0.001 and *β*_graph_ = 0.1, and loss balancing hyperparameters *λ*_data_ = 1 and *λ*_graph_ = 0.2. Model optimization was performed using the RMSProp optimizer with a learning rate of 2 × 10^=>^ and a batch size of 128. Cells were split into training and validation sets using a 90/10 split. The total validation loss was used for early stopping, and the model with the lowest validation loss was used for downstream analysis.

### Sampling Outputs from MeRN

We rely on three key outputs from MeRN to describe a cell’s metabolic and non-metabolic variability based on input scRNA-seq data. We extract a cell’s metabolic and non-metabolic embeddings as the expected values (means) of the learned variational posteriors, *q*(*m* ∣ *x*) and *q*(*b* ∣ *x*), respectively. These latent representations of cells can be used for common downstream analysis tasks such as clustering, pseudotime inference, and visualization.

### Monte Carlo Decoding

To estimate a cell’s expected reaction activity, which corresponds to the reaction activations *r*, we compute the average over 8 Monte Carlo samples drawn with independent random seeds. We selected this number based on the average reaction correlation across cells between independently sampled reaction activations. Specifically, as we increased the number of samples averaged for *r*, the reaction correlation between independently sampled estimates across cells plateaued around 8 samples (**Figure S1I**). Mathematically, letting *K* = 8 denote the number of Monte Carlo samples, we define this expected activity as:

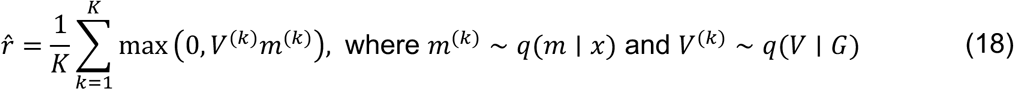

### Ensemble Decoding Across Model Replicates

We also found it beneficial to train multiple replicates of MeRN based on different weight initializations and data splits. A similar experiment comparing reaction correlations between averages of independently trained replicates showed that correlation increased with the number of replicates and plateaued around 5 replicates (**Figure S1J**). We trained 15 independent MeRN replicates for the mouse intestine and 10 replicates for each mouse embryogenesis condition, the mouse cytokine dictionary, the human T cell atlas, and the human NSCLC immune cell atlas. We then used the final ensemble average of r̂ across all replicates for the downstream analysis of reaction activity.

### Hyperparameter Optimization

Hyperparameter selection for MeRN presents a distinct problem because we aim to optimize the model for interpretation rather than simply minimizing validation loss. As a result, we chose to optimize model interpretation consistency across independent training runs.

Given two independently trained models, *X* and *Y*, we assess their consistency using the following procedure:

1. **Cell Latent Space Consistency:** For both the metabolic (*M*) and non-metabolic (*B*) latent spaces, we calculated each cell’s 100 nearest neighbors based on Euclidean distance. We then computed the Jaccard index between each cell’s neighbors from model *X* and model *Y*. To compute a metric of cell latent space consistency between the models, we averaged the Jaccard index across all cells.
2. **Graph Embedding Consistency:** We performed the same nearest-neighbor procedure for the graph embedding (*V*), calculating the Jaccard index using the 25 nearest-neighbor reactions and averaging over all reactions.
3. **Reaction Activity Consistency:** To assess the consistency of the expected reaction activity, we calculated the Spearman correlation for each reaction between the estimated reaction activities, *r̂_X_* and r̂_F_, from models *X* and *Y*, respectively. We averaged these correlations across all reactions.

We performed a hyperparameter grid search on data from the small intestine to select the latent dimension sizes. During the dimension sweep, *z* = *u* = *d*, where *d* ∈ {8,15,25,35,45,55,75}. We also varied the KL divergence weights over *β*_cell_ ∈ {0.0001,0.001,0.005,0.01} and *β*_graph_ ∈ {0.01,0.1}. We trained 15 model replicates for each hyperparameter configuration, resulting in a total of 840 models. Trends across latent dimension sizes were consistent across KL loss weight choices, with consistency metrics plateauing around metabolic dimension 25 or 35 and non-metabolic latent space consistency peaking at dimension 15 (**Figures S1A-D**). We selected the non-metabolic latent size to be *u* = 15 and the metabolic latent size to be *z* = 25 as reasonable choices to maximize our consistency metrics while reducing model size.

To select KL weights, we further inspected consistency dynamics for models trained with *z* = 25 and *u* = 15 on the intestine, cytokine dictionary, and WT embryogenesis datasets. We observed reaction consistency decrease when *β*_cell_ increased past 0.001. Graph, metabolic, and non-metabolic latent space consistency remained relatively stable. Furthermore, *β*_graph_ = 0.1 generally produced higher consistency across all metrics except non-metabolic latent space consistency (**Figures S1E-H**). We selected a cell KL weight of *β*_cell_ = 0.001 and a graph KL weight of *β*_graph_ = 0.1, because this combination optimized reaction activity consistency while regularizing the cell latent spaces as much as possible.

### Accounting for Batch Effects

Inspired by scVI^32^, we account for technical batch effects and other data covariates through the design of our flexible decoders. Specifically, we concatenate a one-hot-encoded batch vector, *c*, to both the reaction activations *r* and the non-metabolic latent representation *b*. By passing these concatenated representations through the decoders, we allow the respective weight matrices, *W*_met_ and *W*_back_, to learn batch-specific additive effects. This design allows the decoders to account for technical variation without including the batch covariate in the latent representations.

### MeRN Software Package

We implemented MeRN as a Python package available at https://github.com/wagnerlab-berkeley/mern. The package contains code for training MeRN, key analysis functions such as DDP calculation, and visualization tools. All models in this paper were trained with v1.0.1 except for the embryogenesis models, which were trained with v1.0.0. The key difference between these versions was a bug that included five additional self-loops in the metabolic guidance graph in v1.0.0. MeRN replicates trained on the small intestine data with v1.0.0 and v1.0.1 were indistinguishable according to the consistency metrics described above.

### DDP Construction and Evaluation

#### Topology-Constrained DDP Construction and Linkage Matrix

DDPs are calculated using topology-constrained agglomerative clustering with complete linkage. For a selected set of cells, let ℛ denote the set of reaction nodes and let ***S*** = [*s_i_*_+_] denote the reaction correlation matrix, where

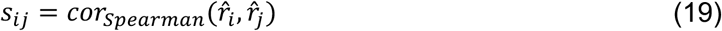

is the Spearman correlation between the activities of reactions *i* and *j* across the selected cells. The clustering procedure returns both DDP assignments and a linkage matrix ***L*** that represents the complete merge hierarchy. We construct these outputs using the following iterative procedure:

1. **Initialization**: Each reaction *i* ∈ ℛ is initialized as a singleton cluster, *C_i_* = {*i*}.
2. **Adjacency and Complete Linkage:** Two clusters *C_a_* and *C_b_* are considered adjacent if at least one edge (*u*, *v*) ∈ ℰ connects a reaction *u* ∈ *C_a_* to a reaction *v* ∈ *C_b_*. For adjacent clusters, complete-linkage similarity is defined as the minimum correlation between their constituent reactions:

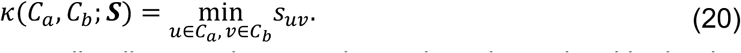

1. **Merge Selection:** Among all adjacent cluster pairs, select the pair with the largest complete-linkage similarity, *κ*^∗^.
2. **DDP Extraction:** At the first iteration for which *κ*^∗^ is strictly less than the correlation threshold *τ*, save the current clusters before performing that merge as the candidate DDPs. The clustering then continues so that the full linkage history is retained. If no proposed merge crosses *τ*, the terminal clusters are used as the candidates.
3. **Merging:** Merge *C_a_* and *C_b_* at linkage distance 1 − *κ*^∗^ and replace them with *C_a_* ∪ *C_b_*.
4. **Termination and Completion:** Repeat steps 2–5 until no graph-supported merge remains. Remaining disconnected roots are joined only to complete ***L***; these completion merges do not alter the saved DDP assignments.
5. **Size Filtering:** Retain candidate DDPs containing at least the minimum number of reactions *d* and assign the retained clusters consecutive DDP identifiers.

The cell set used to calculate ***S*** can be restricted for a particular analysis. Unless otherwise noted, we used *τ* = 0.7 and *d* = 3 for analyses in this paper. We depict this process in Figure 3A.

### Weakest Link Analysis

The weakest link analysis identifies structural breaks in DDPs defined in a reference condition when the same reactions are evaluated in an alternate condition, such as a gene knockout, disease state, or drug perturbation. Let *q* ∈ {*ref*, *alt*} index the two conditions. For each condition, we calculate a reaction correlation matrix ***S***^(*q*)^ as defined above and run the same topology-constrained clustering procedure to obtain a linkage matrix ***L***^(*q*)^. DDP membership is defined only from ***S***^(*ref*)^ and is held fixed when scoring the alternate condition.

We convert each linkage matrix into a cophenetic correlation matrix ***H***^(*q*)^ = [ℎ^(*q*)^]. For reactions *i* and *j*, let *d*^(*q*)^ be the cophenetic distance—the linkage distance at which the two reactions first belong to the same cluster in ***L***^(*q*)^. We express this distance on the correlation scale as

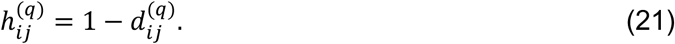

Thus, ***S***^(*q*)^ measures the direct pairwise correlation between reaction activities, whereas ***H***^(*q*)^ describes the correlation level at which each reaction pair becomes connected in the complete topology-constrained hierarchy.

For a reference DDP *D*, let *P*(*D*) denote all unordered pairs of distinct reactions in *D*. We define the minimum raw and cophenetic correlations in condition *q* as

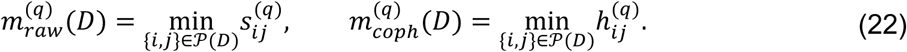

The two weakest-link statistics are the decreases from the reference to the alternate condition:

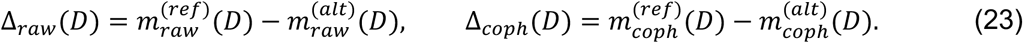

Positive values indicate that the weakest within-DDP relationship is lower in the alternate condition.

### Pathway Scoring

MeRN’s expected reaction activity estimates (*r̂*) allow the scoring of arbitrary groupings of reactions. This is particularly useful for evaluating the activity of predefined biological pathways (e.g., KEGG pathways) or our derived DDPs. To estimate the overall activity of a given pathway, we compute the average expected activity across all of its component reactions as follows:

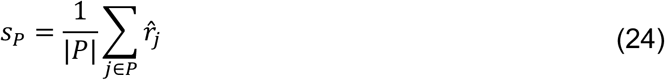

where *s_P_* represents the activity score for a pathway *P*, |*P*| is the total number of reactions contained within the pathway, and *r̂_j_* denotes the expected reaction activity for reaction *j* ∈ *P*.

### Metabolic Directions

MeRN-based metabolic directions enable interpretable analysis of metabolic perturbation effects. Given a reference condition *ref* and an alternative condition *alt*, let *m̄_ref_* and *m̄_alt_* denote the mean metabolic latent positions of their cells. We define the metabolic direction vector as

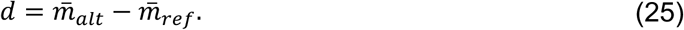

To interpret the reaction activity changes associated with this direction, we project *d* through the reaction embedding matrix:

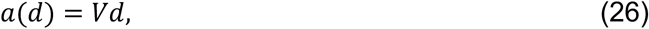

where *a_j_*(*d*) is the signed loading of reaction *j* along the metabolic direction. For a DDP *D*, we define its predicted change along *d* as the average reaction loading,

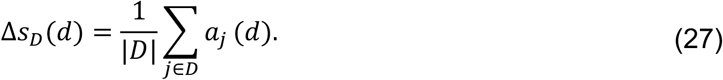

### Metabolic Pathway Visualizations

To visualize metabolic reaction connectivity in context, we used the *custom_pathway_plot* and *kegg_pathway_plot* functions from the MeRN Python package. We constructed these graph layouts based on the KEGG KGML used to construct the MeRN reaction-reaction graph, including edges between metabolites that were substrate-product pairs for a KEGG reaction entry. Metabolite node coordinates were selected using the *graphviz_layout* function from *NetworkX* for custom layouts or based on the metabolite coordinates in the KEGG pathway plot. We note the metabolic pathway visualizations are visualization tools, not perfect representations of the underlying metabolic topology.

### Dataset-Specific Downstream Analyses Small Intestine (Figure 2)

#### Alternative metabolic inference methods for benchmarking

We benchmarked MeRN against a variety of methods for inferring metabolic state from transcriptomic data. Generally, these methods take transcriptomic data as input and return cell-level reaction activity estimates. Although the output reactions generally represent similar underlying reaction sets, these methods vary in their choice of genome-scale metabolic models (GEMs), necessitating the harmonization of reaction identifiers across GEMs for comparable benchmarking. To map all method outputs to the set of KEGG global pathway map reaction entries, we used MetaNetX version 4.5^86^. We first mapped the reaction IDs from each reaction entry node to a global MetaNetX identifier. We then matched this identifier with reaction IDs from other GEMs. For MetroSCREEN, we additionally used KEGG reaction identifiers in the Human-GEM MIRIAM annotations. For the minority of KEGG reaction nodes that mapped to multiple reaction estimates from a given method, we calculated all pairwise Spearman correlations across cells. If every pairwise correlation exceeded 0.9, we randomly selected one reaction using a fixed seed. Otherwise, we retained the estimate only when exactly one matched reaction was annotated as cytosolic, excluding reaction nodes with zero or multiple cytosolic matches.

### Compass

Compass v1.0.0^23^ was run on log-normalized counts from the preprocessed small intestine data. Cell-specific reaction penalties were converted to reaction consistencies and used for downstream benchmarking. A total of 298 KEGG reaction nodes were matched to Compass outputs.

### scCellFie

scCellFie v0.5.0^28^ was run on raw counts from the preprocessed small intestine data using parameters from the mouse brain tutorial. Reaction activities were used for reaction-level analyses and metabolic task scores for comparison with MeRN in Figure 2G. A total of 202 KEGG reaction nodes were matched to scCellFie reaction activities.

### MetroSCREEN

Due to incompatibilities between the updated GSVA R package^87^ and the MetroSCREEN package^31^, we manually implemented the ‘cal_MetaModule’ function with the same GSVA parameters as the original function. We used the MetroSCREEN-provided metamodules and gene-protein-reaction (GPR) rules. We also humanized the mouse genes from the small intestine dataset because no mouse metamodule version was provided. A total of 436 KEGG reaction nodes were matched and estimated based on MetroSCREEN metamodule activities.

### KEGG MetroSCREEN

To bypass the mapping step between alternate GEMs and KEGG reactions, we adapted the MetroSCREEN method to KEGG by using the KEGG reaction-to-gene mapping for mouse. Importantly, GPR rules were omitted from this adaptation. This direct mapping yielded estimates for 1,023 KEGG reaction nodes.

### scFEA

scFEA v1.1-40-g4c1fb76^24^ was run on both raw and log-normalized counts for enterocytes from the preprocessed small intestine data following the steps in the tutorial. Estimated module flux was highly correlated between the two, and we selected the flux estimated from the normalized counts for the analysis in Figure 2F. We mapped KEGG reaction nodes used in MeRN to scFEA modules based on the “Module_reaction” sheet of scFEA Supplementary Table S1.

### GSEA and overrepresentation analysis across proximal–distal domains

We computed GSEA normalized enrichment scores (NESs) and overrepresentation combined scores for KEGG 2019 mouse pathways with the prerank and enrichr methods from the ‘gseapy’ package (v1.2.1)^88^, respectively. To simulate a realistic analysis comparing metabolic pathway usage across the proximal-distal domain, we performed one-versus-rest differential expression with a Wilcoxon test for each domain. We passed the signed, standardized Wilcoxon rank-sum test statistics to the prerank function and all genes with a positive log-fold change and adjusted p-value less than 0.001 to the enrichr function, resulting in NES and combined scores for each domain and KEGG pathway.

### Villus region mapping

To map each enterocyte from the Zwick et al. dataset to a villus region, we first used a sample-level one-versus-rest Wilcoxon test to identify differentially expressed genes for each villus region from the Harnik et al. bulk RNA-seq data^20^. We constructed a gene set for each region from the genes with a signed, standardized Wilcoxon rank-sum test statistic greater than 2, resulting in 510, 196, 132, 40, and 623 genes for each region. We then used the Scanpy score_genes function to score enterocytes based on the genes for each villus region. We classified enterocytes as villus-bottom landmarks when their V1 score was above the 85th percentile and their V6 score was below the 85th percentile. We calculated pseudotime from the cell closest to the median metabolic embedding of these landmark enterocytes. We then scaled the villus region gene scores over the enterocytes. Finally, we used a Gaussian filter with *σ* = 50 to smooth the scaled region scores over enterocytes ordered by their pseudotime and assigned each enterocyte to a villus region based on the maximum smoothed and scaled score at its pseudotime position. This procedure resulted in no enterocytes assigned to V5, so we excluded V5 from the remaining analyses.

### Preprocessing of protein data

We downloaded the protein data from Harnik et al. and applied their reported filtering criteria to the supplied combined protein table. Specifically, we retained proteins whose mean sum-normalized iBAQ (intensity-based absolute quantification) fraction across regions exceeded 10^=Y^ in at least one mouse and whose mouse-level mean was nonzero in at least two mice. The mean iBAQ fraction for V1 was calculated from three mice because the M3 V1 measurement was absent from the combined table. Unless otherwise noted, we used the mean iBAQ fraction over all provided mice in each region.

### Reaction profiles across villus zones

For each modality and metabolic inference method, we estimated the activity of each reaction in ℛ over the regions of the villus. For the scRNA-seq data, we summed the counts of each gene across enterocytes in each region and normalized across genes within each region. For bulk and single-cell RNA-seq, we estimated each reaction in ℛ as the maximum of any matched gene based on our reaction–gene mapping. For protein, we took the mean abundance over genes matching each reaction. Finally, for MeRN, Compass, scCellFie, MetroSCREEN, and KEGG MetroSCREEN, we took the mean reaction activity over all enterocytes in each region.

### Selecting reactions for benchmarking in **Figure 2H**

We selected zonated reactions over the villus with a mouse-level ANOVA test over protein estimates of reaction activity. We retained all 395 reactions with an FDR < 0.25 after Benjamini-Hochberg correction. For each pairwise comparison between methods or modalities, we computed the Spearman correlation over the set of common reactions.

### DDPs and Embryogenesis (Figure 3)

#### Cell selection for DDP construction

A minimum correlation threshold of 0.7 and minimum DDP size of 3 were used to calculate all DDPs. The small intestine enterocyte and stem cell DDPs used all 5,383 and 6,239 enterocytes and stem cells, respectively. The embryogenesis DDPs were calculated on all 11,431 WT MS2 cells (MS2 assignment is described in the accompanying manuscript by Dias et al.).

### Variance explained by DDP and KEGG pathway scores

For each reaction, we fit an ordinary least squares model predicting reaction activity from its assigned DDP score or KEGG pathway score(s) in the same cells. We recorded the *R*^2^ for each reaction and calculated the mean *R*^2^ across reactions covered by each pathway definition. We filtered out KEGG’s global and overview pathways. We calculated coverage fraction as the fraction of reactions included in a DDP or KEGG pathway. We calculated DDPs at minimum correlation thresholds ranging from 0.05 to 0.90 using a minimum DDP size of 3.

### Cell-cycle position and phase assignment with Tricycle

We estimated cell cycle positions using the R *tricycle* package (v1.6.0)^89^, applied separately to the wild-type and Δ*Folr1* scRNA-seq data using the *Runtricycle* function from *SeuratWrappers* (v0.3.5), specifying the *species* parameter as “mouse” with all other defaults applied. Cell cycle phases were assigned based on position, where θ ∈ (0,2π), using the following thresholds: *S* (0.5π ≤ θ < π), *G2* (π ≤ θ < 1.5π), *M* (1.5π ≤ θ < 1.75π), and *G0/G1* (θ ≥ 1.75π or θ < 0.5π).

### Cytokine Dictionary (Figure 4)

#### Ordinary least squares model of DDP activity

To assess how cell type identity or cytokine treatment explained metabolic DDP activity, we used a fixed-effects model of DDP activity. We calculated DDPs from a fixed 10% sample of 38,670 cells with a minimum correlation threshold of 0.7 and a minimum DDP size of three. We then averaged DDP activity for cell type-treatment-mouse combinations, excluding combinations with 25 or fewer cells. For each DDP, we standardized the activity over all remaining groups and fit an ordinary least-squares model that included mouse, sum-coded cell type, and PBS-referenced cytokine treatment as covariates, along with a cell type-treatment interaction term.

### Metabolic directions analysis of cytokine perturbation in T cells

To select cytokines with consistent cell type-specific changes across biological replicates, we calculated metabolic directions (described above) between cytokine-perturbed cells for each replicate and the average location of PBS (control) cells for CD4+ and CD8+ T cells separately. For each cell type, we retained cytokines with an average pairwise cosine distance of less than 0.5 for further analysis, keeping 12 cytokines for CD4+ T cells and 15 cytokines for CD8+ T cells. For each retained cell type-cytokine pair, we recalculated the metabolic direction based on the average location of perturbed cells across all three biological replicates. We performed the same procedure using non-metabolic embeddings to calculate non-metabolic direction vectors for each cell type-cytokine pair. To calculate the DDP activity changes in Figure 4F, we averaged the predicted reaction activity changes across reactions in each DDP as described above.

### Differential-expression analysis of cytokine responses

For the CD4+ T cell differential expression analysis, we conducted cell-level Wilcoxon tests comparing each cytokine treatment with PBS. For the CD8+ T cell differential expression analysis, we pooled cells treated with IL-2, IL-18, or IL-21 and performed a cell-level Wilcoxon test against PBS-treated cells.

### Pan-Cancer T Cell Atlas (Figure 5)

#### Linear mixed-effects model of DDP activity

Atlas DDPs were constructed from a fixed 5% sample of 18,669 cells with a minimum correlation of 0.7 and a minimum size of 3. Scaled DDP scores were averaged by patient, tissue location, and T cell subtype, excluding combinations with 25 or fewer cells. We then fit a linear mixed-effects model for each DDP with T cell subtype, tissue location, and cancer type as fixed effects, along with a patient random intercept. T cell subtype coefficients are relative to naïve CD8+ T cells (CD8.c01.Tn.MAL), tissue location coefficients are relative to peripheral blood (‘P’), and cancer type coefficients are relative to melanoma.

### DDP trajectory construction

To identify DDPs associated with stages of T cell function, we represented each DDP by its linear mixed-effects model coefficients across six ordered CD8+ T cell states along the exhaustion trajectory defined by Zheng et al.^68^ We performed Ward hierarchical clustering using Euclidean distances between these coefficient profiles and cut the dendrogram into five clusters. We named the clusters based on the coefficient patterns shown in Figure 5A.

### Determining Treg-specific DDPs in **Figure 5D**

We conducted three one-sided Wald tests for each DDP to test for higher Treg activity (*β*_Treg_ − *β*_Tex_ > 0), strong Treg association (*β*_Treg_ > 0.35), and weak TEX association (*β*_Tex_ < 0.45) with Benjamini-Hochberg correction for each test family. Treg-specific DDPs were DDPs with FDR < 0.05 for all three tests.

### NSCLC Immunotherapy (Figure 5)

#### Calculating AUCs in Figure S5E

The responder label was assigned to patients with a major pathological response (MPR) or pathological complete response (pCR), and the non-responder label to the remaining patients. For each cell type, we retained patients with more than 25 cells. Cell types were included if at least 25 responders and 25 non-responders met this threshold out of a total of 130 responders and 112 non-responders after excluding one patient with an unknown response. We fit separate L2-regularized logistic regression models using either patient-averaged DDP scores or patient-averaged non-metabolic embedding coordinates. Predictors were standardized within each training fold, and models used class balancing and five-fold stratified cross-validation. We calculated the mean ROC AUC across folds.

### Response-associated DDP analysis

We averaged DDP activity by patient and cell type, retaining combinations with more than 25 cells. For each DDP in each cell type represented by at least 25 responders and 25 non-responders, we compared patient-level scores using a two-sided Mann-Whitney U test. We calculated Cohen’s *d* as the responder mean minus the non-responder mean, divided by the pooled standard deviation. P-values were adjusted across DDPs within each cell type using the Benjamini–Hochberg procedure.

### Curating a gene set of Treg activation

We constructed the activated Treg gene set from Alvisi et al.^90^ by intersecting CCR8+ICOS+ versus CCR8-ICOS-differentially expressed genes with the human-mouse conserved tumor-Treg genes, retaining genes with positive log2 fold changes, and selecting the top 49 by log2 fold change. We selected 49 genes to match the Gavish et al.^91^ gene signatures.

---

**Supplementary Figure 1.**
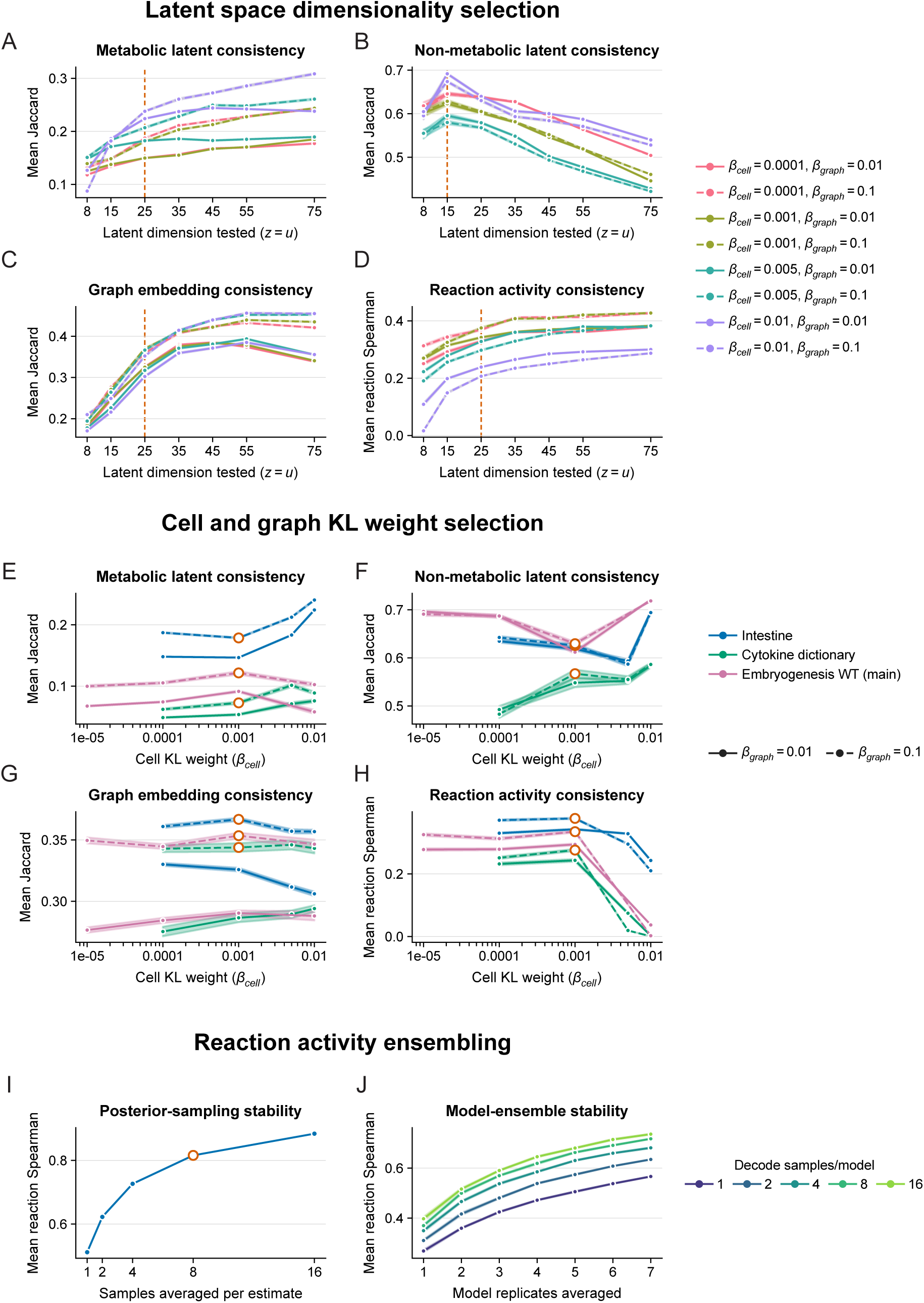
We constructed three metrics to assess the consistency of MeRN’s learned outputs between two independently trained replicates: 1. Latent space consistency (metabolic and non-metabolic): the average Jaccard similarity between the 100-nearest-neighbor sets for each shared cell in the two replicates, averaged across cells. 2. Graph embedding consistency: the average Jaccard similarity between the 25-nearest-neighbor sets for each reaction in the two graph embeddings, averaged across reactions. 3. Reaction activity consistency: the mean per-reaction Spearman correlation between reaction activities predicted by each model. **(A–D)** To select latent dimension sizes, 15 MeRN models were trained on the small intestine data at each tested shared latent dimension (*z* = *u*), cell KL weight (*β*_cell_), and graph KL weight (*β*_graph_). The average metabolic latent space consistency **(A)**, non-metabolic latent space consistency **(B)**, graph embedding consistency **(C)**, and reaction activity consistency **(D)** were calculated across all pairwise model-replicate comparisons. The dashed orange lines show the selected dimensions: *z* = 25 for A, C, and D, and *u* = 15 for B. Color denotes *β*_cell_, line style denotes *β*_graph_, and shaded ribbons denote the standard error. **(E–H)** 15, 10, and 10 MeRN replicates were trained on the small intestine, cytokine dictionary, and mouse embryogenesis datasets, respectively, with metabolic latent dimension *z* = 25 and non-metabolic latent dimension *u* = 15, across the tested *β*_cell_ and *β*_graph_ weights. The average metabolic latent space consistency **(E)**, non-metabolic latent space consistency **(F)**, graph embedding consistency **(G)**, and reaction activity consistency **(H)** were calculated across all available pairwise model-replicate comparisons for each dataset. Orange open points denote the selected configuration (*β*_cell_ = 0.001, *β*_graph_ = 0.1). Color denotes dataset, line style denotes *β*_graph_, and shaded ribbons denote the standard error. **(I)** For each number of samples averaged per estimate, two independent reaction-activity estimates were generated from each small intestine model by averaging that many independent decodes per estimate. Fifteen random split comparisons were performed for each model and sample-size value. The mean per-reaction Spearman correlation between the two estimates was calculated. The orange open point denotes the selected eight samples per reaction-activity estimate. **(J)** Same as (I), except reaction-activity estimates were generated by averaging model replicates, using the specified number of model replicates per side and decodes per model. Fifteen random split comparisons were performed for each setting. Color denotes the number of decodes averaged per model.

**Supplementary Figure 2.**
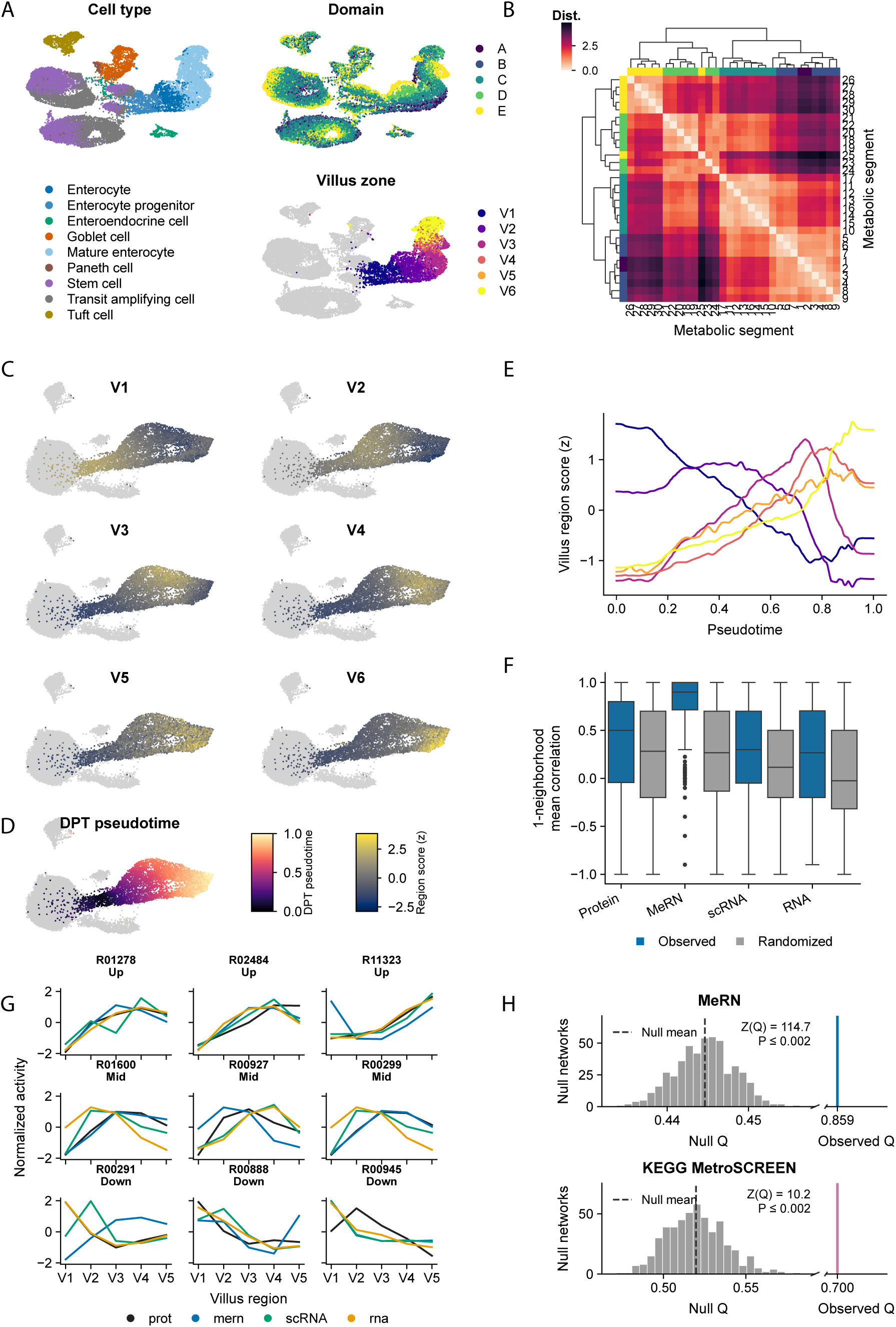
**(A)** UMAP representations of scRNA-seq data from the non-metabolic latent space, colored by cell type, proximal-distal domain, and villus zone. **(B)** Pairwise Euclidean distances between the mean metabolic latent space coordinates of enterocytes from each proximal-distal segment. Segments were ordered by hierarchical clustering of the distance matrix. Row and column colors denote domain assignments. **(C)** A gene set for each villus zone was defined as the genes with a score greater than 2 in a one-versus-rest Wilcoxon differential-expression test of bulk RNA-seq data from Harnik et al.^20^ Enterocytes were scored for each gene set, and the resulting z scores were visualized on the metabolic UMAP. **(D)** Enterocytes with V1 scores above the 85th percentile and V6 scores below the 85th percentile were classified as bottom cells. The bottom cell closest to the median metabolic embedding of this group was selected as the root, and diffusion pseudotime (DPT) was computed on the metabolic latent space and visualized on the metabolic latent UMAP. **(E)** Z-scored villus-zone scores, smoothed with a Gaussian filter (sigma = 50 cells), for enterocytes ordered by DPT pseudotime. Enterocytes were assigned to the villus zone with the maximum smoothed scaled score. **(F)** For each reaction, the mean Spearman correlation between itself and its immediate neighbors in the metabolic topology network was computed across the five villus zones for each modality, excluding reaction pairs with overlapping enzyme-gene sets. Blue denotes observed distributions; gray denotes distributions from 10 random relabelings of graph nodes. **(G)** Enterocyte reaction scores were averaged within each villus region and standardized separately for each method or modality. Three reactions each classified as increasing, decreasing, or centrally peaking by their protein profile were randomly selected with a fixed seed for visualization. **(H)** For the MeRN and MetroSCREEN reaction networks in Figure 2I,J, unweighted Newman modularity^56^ was computed on the largest connected component. Null distributions were generated from 500 connected, degree-preserving edge-rewired networks, each with 10 successful swaps per edge. Colored bars denote observed modularity, and gray histograms denote null modularity distributions.

**Supplementary Figure 3.**
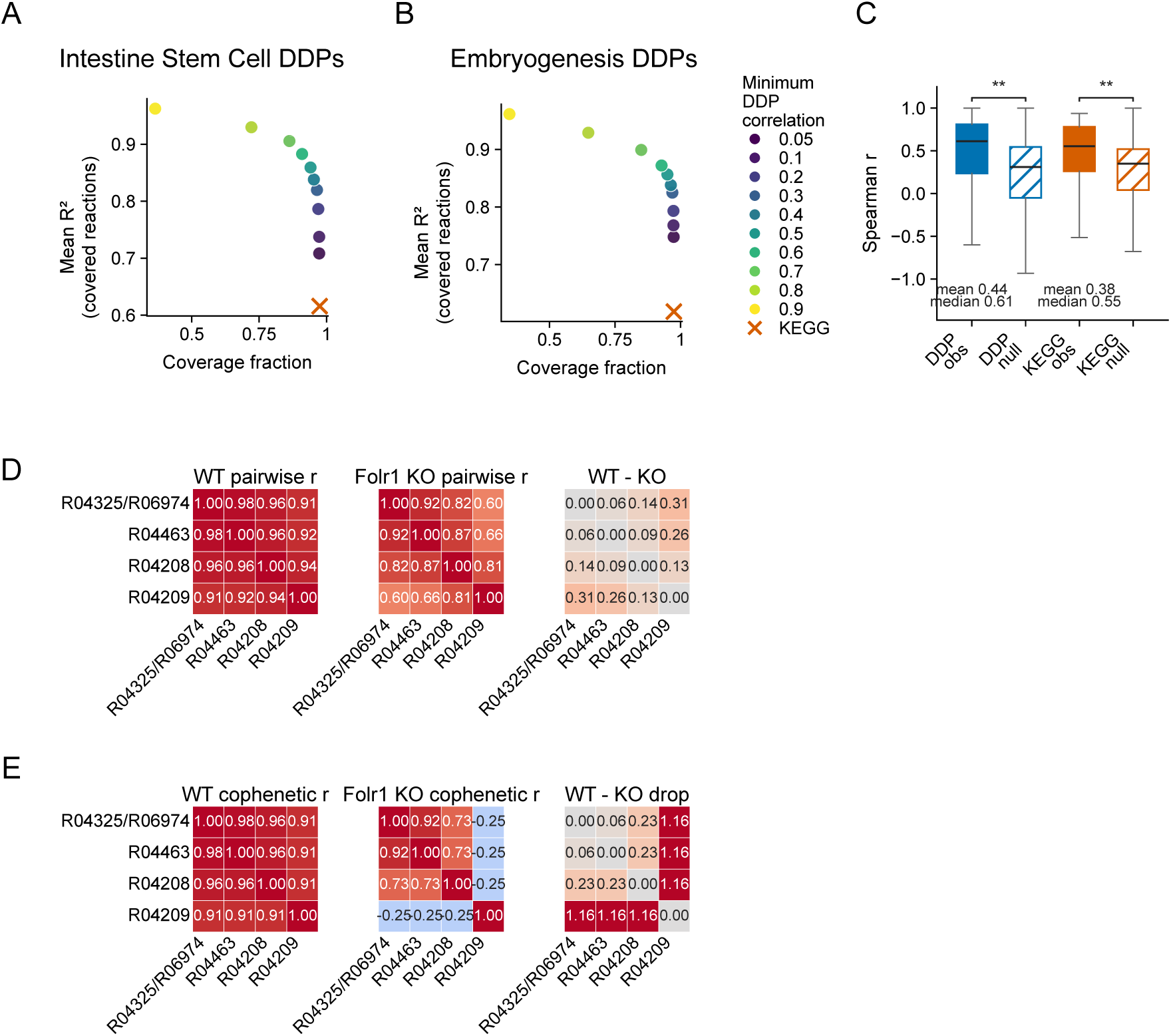
**(A)** DDPs were computed at ten minimum-correlation thresholds from 0.05 to 0.90 in intestinal stem cells. For each threshold, coverage was the fraction of all reactions assigned to a DDP, and mean R² was calculated across covered reactions from in-sample OLS models predicting each reaction’s activity from its DDP score. The corresponding KEGG value, obtained using all assigned KEGG pathway scores as predictors, is shown in orange. **(B)** Same as panel (A), except for WT MS2 embryogenesis cells (defined in the accompanying manuscript by Dias et al.). **(C)** For each reaction with a zonated protein profile, the mean Spearman correlation with other reactions in the same DDP or KEGG pathway was calculated across five villus zones (V1-V4 and V6), excluding reaction pairs with overlapping enzyme-gene sets. Observed distributions were compared with 100 random membership nulls that preserve DDP group sizes or KEGG pathway sizes and reaction membership degrees. Solid boxes denote observed values, and hatched boxes denote null values. Brackets report empirical one-sided tests of the observed mean against the corresponding null distribution. **(D)** Pairwise Spearman correlations among reactions in DDP 44 across WT and Folr1 KO MS2 cells, together with the WT-minus-KO difference. **(E)** Same as panel (D), except for pairwise cophenetic correlations.

**Supplementary Figure 4.**
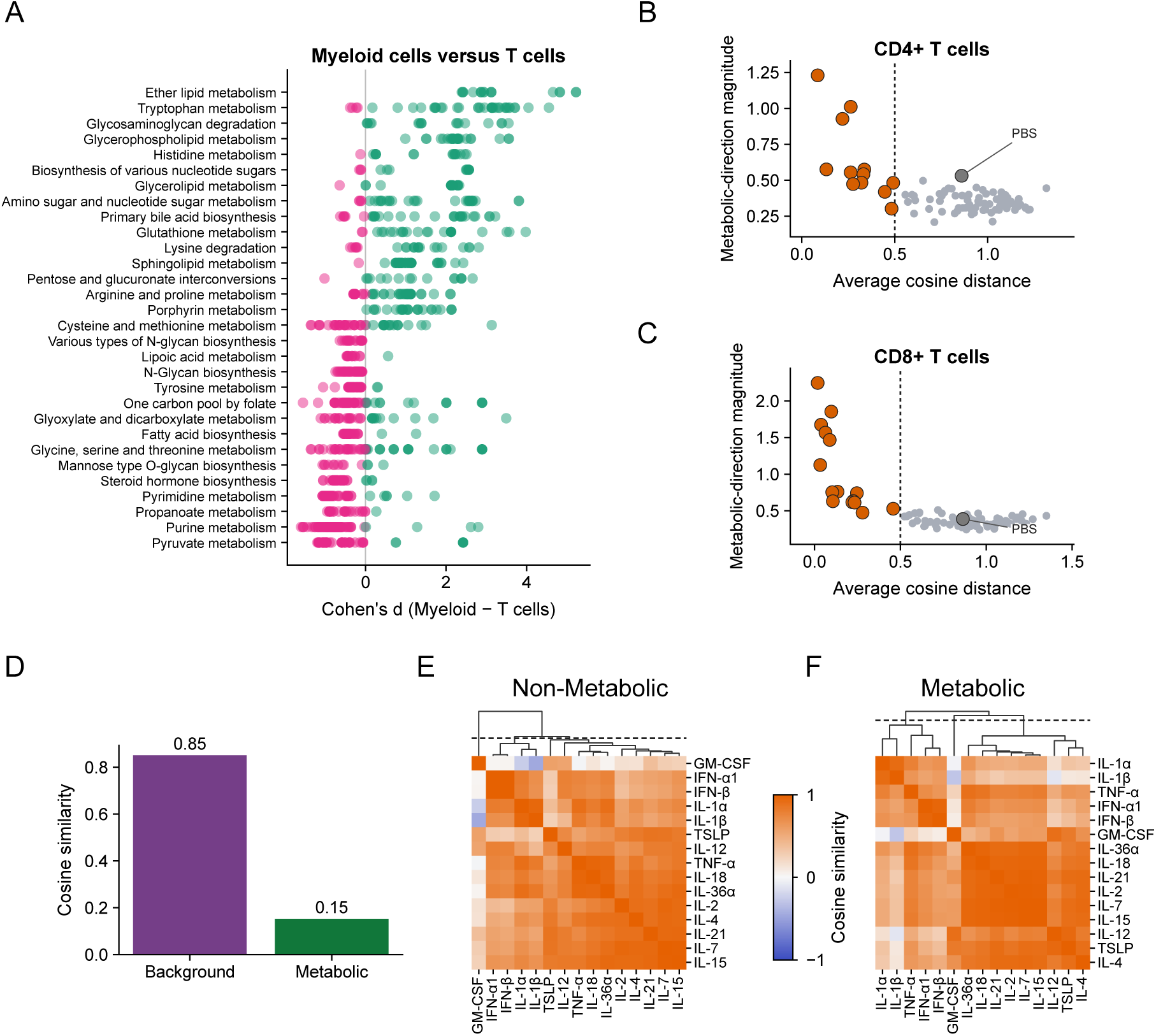
**(A)** Reaction-level Cohen’s d values comparing myeloid cells with T cells are shown for 15 KEGG pathways at each extreme of the pathway-level Cohen’s d ranking; eligible pathways contained more than 10 reactions and were represented in the mouse KEGG resource. **(B) F**or each cytokine in CD4+ T cells, the mean of the three pairwise cosine distances among metabolic direction vectors from three biological replicates is plotted against the mean direction magnitude. Cytokines below the cosine distance threshold of 0.5 (denoted by a vertical dotted line) are colored orange and kept for downstream analysis. The negative control (PBS) is highlighted in grey. **(C)** Same as panel (B), except for CD8+ T cells. **(D)** Cosine similarity between the IL-36α and IL-1β non-metabolic and metabolic direction vectors in CD4+ T cells. **(E)** Cosine similarity between non-metabolic direction vectors for CD8+ T cells. Cytokines were ordered by average-linkage hierarchical clustering of the similarity matrix. The dashed line marks the dendrogram height that produces three clusters. **(F)** Same as panel (E), except for metabolic direction vectors.

**Supplementary Figure 5.**
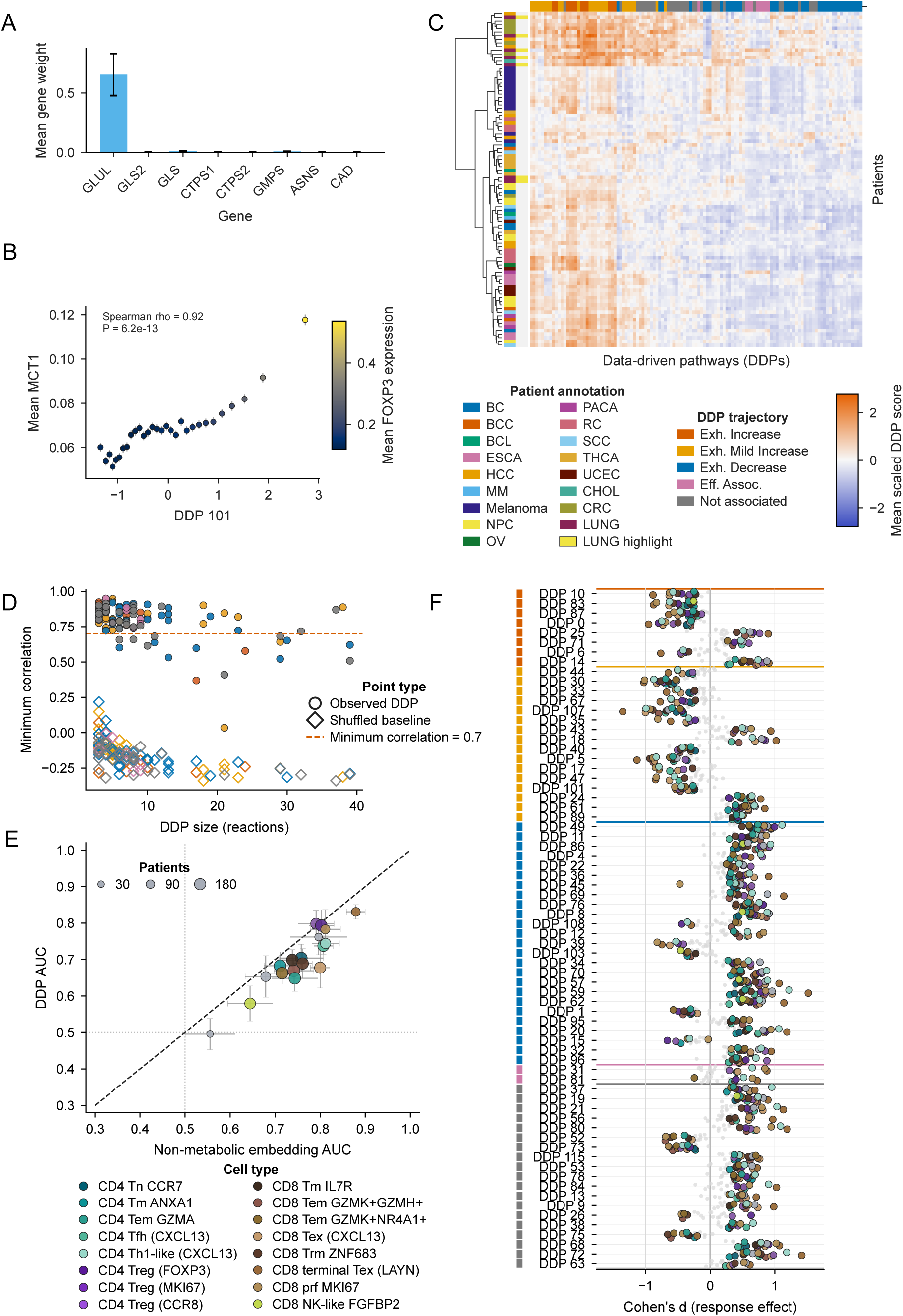
**(A)** Mean gene weights for the glutaminolysis reaction group R00253 R00256 (GLUL, GLS2, GLS, CTPS1, CTPS2, GMPS, ASNS, and CAD) across ten MeRN replicates; error bars show the standard error across replicates. **(B)** Mean MCT1/SLC16A1 expression versus mean DDP 101 activity in 30 quantile bins across cells in the pan-cancer atlas. Color denotes mean FOXP3 expression, and error bars show the standard error of MCT1/SLC16A1 expression within each bin. **(C)** Mean scaled DDP activity by patient in tumor-derived CD8 Tex (CXCL13) cells. Patient-cell groups with 25 or fewer cells were excluded. Patients and DDPs were ordered by Ward hierarchical clustering of Euclidean distances and annotated by cancer type, LUNG status, and DDP trajectory group. **(D)** For each atlas-defined DDP, the minimum pairwise Spearman correlation among its reactions was computed across a fixed random sample of 25,000 NSCLC T cells. The shuffled baseline was generated by drawing one random reaction set of the same size as each DDP. Filled circles denote observed DDPs, open diamonds denote shuffled sets, and the dashed line marks a minimum correlation of 0.7 (used for the original DDP construction). Only DDPs with a minimum correlation above 0.7 were kept for downstream analysis. **(E)** DDP activity and the non-metabolic embedding were separately averaged at the patient level within each cell type and used to predict pathological response with L2-penalized logistic regression. Points show mean ROC-AUC from five-fold stratified cross-validation; error bars show the standard error across folds, color denotes cell type, and point area denotes patient count. Patient-cell type groups with 25 or fewer cells were excluded, and each analyzed cell type required at least 25 responders and 25 non-responders. **(F)** Patient-level DDP activity was compared between responders and non-responders within each cell type. Patient-cell type groups with 25 or fewer cells were excluded, and each analyzed cell type required at least 25 responders and 25 non-responders. DDPs significant at Benjamini-Hochberg FDR at or below 0.05 in at least six cell types were retained. Each point represents Cohen’s d for a given DDP and cell type. Colored points indicate FDR-significant effects; smaller gray points are not significant.

